# An archaeal CBASS shows protective activity against both chronic and lytic viruses

**DOI:** 10.1101/2024.09.12.612678

**Authors:** Deepak Kumar Choudhary, Himani Singla, Dana Vassover, Noam Golan, Leah Reshef, Hadar Shukrun, Aime Bienfait Igiraneza, Mechthild Pohlschröder, Uri Gophna

## Abstract

Many cyclic-oligonucleotide-based anti-phage signalling systems (CBASS) defend against viral infections by depleting cellular NAD^+^ levels, eventually leading to dormancy or death. This abortive infection strategy is beneficial in stopping fast lytic infections, as cells die before spreading the virus to neighbouring cells. However, in chronic viral infections, which often occur in archaea, abortive infection could be detrimental, as the cost of immunity may outweigh that of infection. Here we study an archaeal CBASS system (H-CBASS2) that was expressed in the model species *Haloferax volcanii* DS2 and *Haloferax gibbonsii* LR2-5. We show that the system provides protection to *H. gibbonsii* against a lytic tailed haloarchaeal virus HFTV1 by depleting NAD^+^, similar to what has been observed for lytic phages in bacteria. H-CBASS2 is also triggered, though with much slower activity, during infection with the chronic, non-lytic virus HFPV-1, and promotes virus clearance after several passages without killing host cells. Moreover, cells that clear the HFPV-1 infection become substantially more resistant to subsequent infections, due to mutations in envelope-associated proteins. Cell death by NAD^+^ depletion only occurs after a very long infection with HFPV-1 on solid medium. These findings suggest that the magnitude of H-CBASS2 response is somehow tuned to the infection type can benefit the host during non-lytic infections, potentially explaining why such systems are relatively common in archaea.

## Introduction

CBASS protect bacterial populations by killing the cell before the phage reaches maturation by diverse mechanisms, thereby stopping infection spread ^1–6^. These systems share a common ancestry with the cyclic GMP–AMP synthase (cGAS)–STING immune pathway of animals^1,7–9^, and act by sensing a viral infection and producing a cyclic-oligonucleotide signal molecule, which is then sensed by an effector protein that later causes early cell death before the virus completes its replicative cycle. CBASS is fairly common and present in over 10% of bacterial genomes and in 5-10% of archaeal genomes ^1,5,10,11^. One of the most common effector types in CBASS is a TIR-SAVED domain protein that upon sensing the signal molecule depletes cellular NAD^+^ levels, resulting in cell death^12,13^.

Abortive-infection systems, such as CBASS, are beneficial in stopping a rapid lytic infection because cells die before spreading the virus to closely-related sister cells. Importantly, cells at advanced stages of lytic infection have nothing to lose from suicide, since they will soon lyse anyway when the phage completes its lifecycle. Conversely, many archaea, in their natural habitats, are infected by non-lytic viruses that chronically co-exist with their hosts for extended periods.

In such chronic infections, the virus continuously replicates and virions are released from host cells over an extended period without causing immediate cell lysis or death^14^. Under such situations, abortive infection should in principle be harmful because the cost of immunity may be higher than that of infection, yet intriguingly CBASS is about as common in archaea as they are in bacteria.

All previously characterized CBASS systems have been studied in bacteria in the context of lytic phage infection. Here, for the first time, we heterologously expressed an archaeal Type II CBASS system from *Haloferax* strain Atlit 48N (hereafter referred to as H-CBASS2), and examined its function in two haloarchaeal host-virus systems, one lytic and the other chronic. We show that while H-CBASS2 provides some protection against lytic infection, it is also active under chronic infection, but at a much slower rate. Nonetheless, even the slower H-CBASS2 activity observed during chronic infection inhibited host growth, while also changing viral protein expression and eventually cleared the viral infection, by favouring the emergence of virus resistant mutants. This suggests that at least some archaeal CBASS systems can tune their activity to different infections, and thus may provide benefits to their hosts even when chronic viral infections outnumber lytic ones.

## Results and Discussion

### H-CBASS2 senses the lytic virus HFTV1 and provides antiviral defence in Haloferax gibbonsii

Haloferax strain 48N encodes two CBASS gene clusters along with multiple additional defence systems [13], making it suboptimal for characterizing the activity of any individual system. We therefore chose to study its Type II CBASS system that encodes a TIR-SAVED effector (H-CBASS2) in two well established host-virus systems: *H. gibbonsii* LR2-5 and its lytic virus, HFTV1^15^ and *H. volcanii* DS2 and its chronic virus HFPV-1^16^. H-CBASS2 was cloned into the replicating plasmid pTA927 under the control of an inducible promoter and transformed into the two heterologous Haloferax hosts, which do not naturally encode any CBASS system (Fig. 1A). To investigate the immune function of H-CBASS2 against lytic infection, we first performed plaque assays using the lytic virus HFTV1. *H. gibbonsii* cells expressing H-CBASS2 showed approximately a 10-fold reduction in plaque formation compared to control cells carrying an empty vector (Fig. 1B & C). This result indicates that H-CBASS2 provides a measurable level of defence against lytic viral infection. We next performed growth experiments and observed that H-CBASS2-expressing cultures collapsed slightly earlier than control cells upon infection; however, this collapse was not as rapid or pronounced as typically observed for classical abortive infection systems (Fig. 1D).

**Figure 1.**
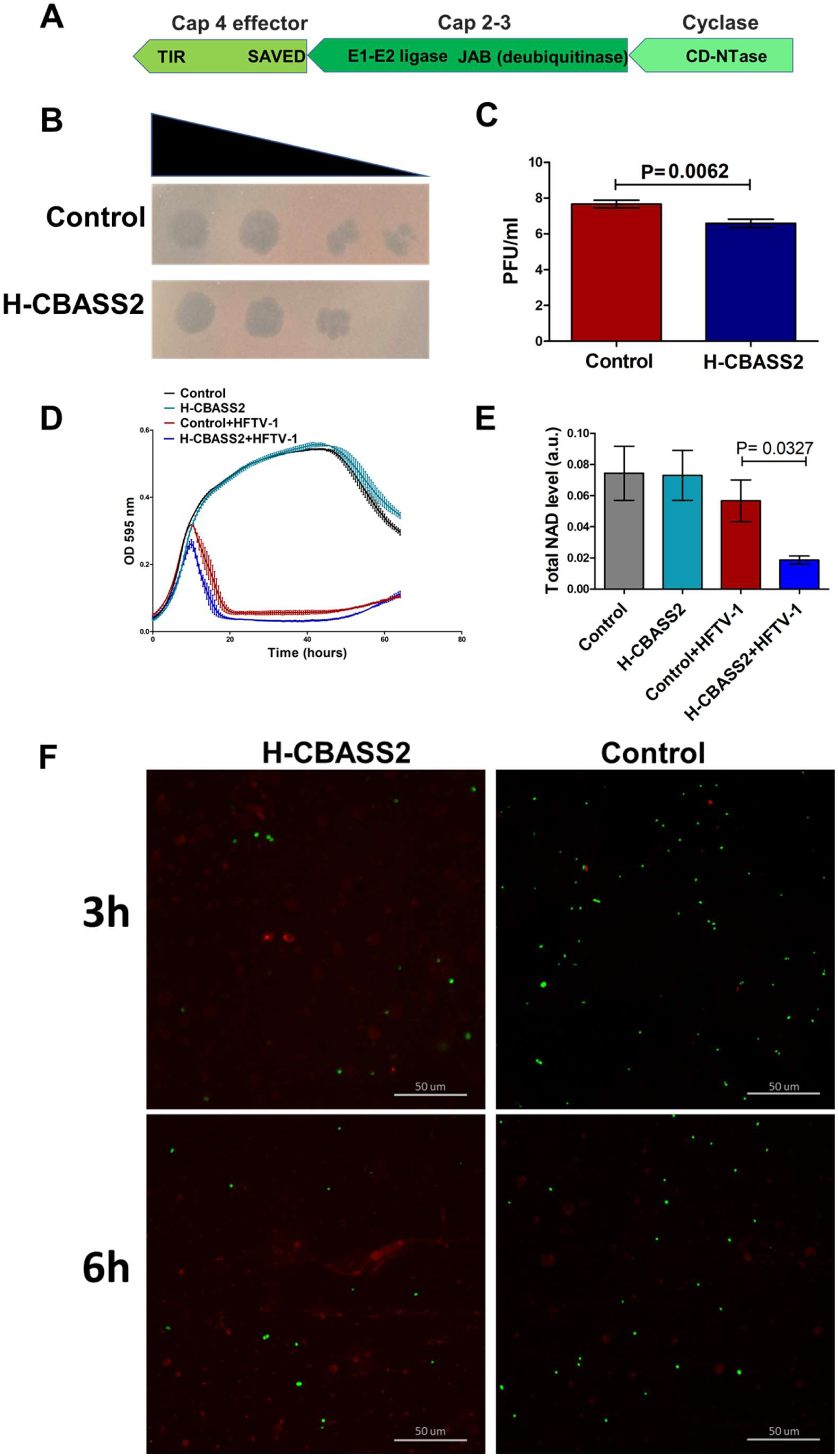
H-CBASS2 detects the lytic virus HFTV1 and confers antiviral protection in *Haloferax gibbonsii*. **A)** The *Haloferax* 48N-CBASS Type II (H-CBASS2) operon. H-CBASS2 genes were cloned into the pTA927 plasmid, under the inducible tryptophanase promoter. The plasmid, referred to henceforth as pCBASS2 was then transformed into uracil auxotroph *H. gibbonsii* LR2-5 **B)** Plaque assay using 10-fold serial dilutions of the virus HFTV-1. H-CBASS2 expressing cells and control cells with an empty vector were mixed with 0.02% top agar and plated onto rich medium. 3 µL of serial dilutions of HFTV-1 were then spotted to quantify plating efficiency. The images are representative of at least three biological replicates. **C)** Plaque assay quantification. Differences among strains were assessed using a paired samples t-test (N = 3)**. D)** Growth experiments were performed with and without HFTV-1 infection. Growth curves are shown for *H. gibbonsii* expressing H-CBASS2 under a tryptophan-inducible promoter (cyan) and the empty vector control (black) in the absence of viral infection. For virus-infected conditions, cells were infected with HFTV-1 at a multiplicity of infection (MOI) of 0.01. Following infection, growth curves are shown for HFTV-1-infected *H. gibbonsii* expressing H-CBASS2 (blue) and the infected control (red). **E)** Total NAD⁺ levels were determined in *H. gibbonsii* strains expressing H-CBASS2, with or without HFTV-1 infection. Cultures were grown to logarithmic phase and subsequently infected with HFTV-1 at a MOI 0.05. Equal cell numbers from each culture were then collected, and total NAD⁺ levels were measured. Data are representative of at least three independent experiments. NAD^+^ values of the different strains were compared using a paired sample t-test (N = 3). **F)** Live/dead assays (green colour represent live cells and red colour represent dead cells) were performed on cells with or without H-CBASS2 expression, followed by infection with HFTV-1 at MOI 0.05. This assay revealed a higher proportion of dead cells in the virus-infected *H. gibbonsii* strain expressing H-CBASS2 compared to the infected control (scale bar: 50 µm). The images are representative of at least three biological replicates.

Nonetheless, Live-Dead staining at 3 h. post-infection showed the presence of much extracellular DNA (diffuse red stain observed outside of cells) and fewer viable intact cells in the H-CBASS2-expressing culture, while the control culture remained intact, as expected for this infection model, where lysis is generally observed only 6h post-infection (Fig. 1F). Thus, the reduction in plaquing efficiency can probably be attributed to premature lysis.

H-CBASS2 has the same architecture as many bacterial type 2 CBASS systems, with effector proteins comprised of a SAVED domain and a TIR domain. In such systems, the SAVED domain senses the cyclic-oligonucleotide signal molecule, while the TIR domain has been shown to be the effector domain and cause cell death by depletion of NAD^+ 12^: H-CBASS2 is therefore expected to function via NAD^+^ depletion. We therefore measured total NAD⁺ levels in intact cells prior to lysis following infection, and observed that indeed infected H-CBASS2-expressing cells exhibited significant depletion of NAD compared to infected control cells (Fig. 1E), while uninfected cells expressing H-CBASS2 did not show any detectable reduction in total NAD levels.

### Expression of *H-CBASS2* in H. volcanii leads to growth delay during chronic virus infection, in a cyclase-dependent manner

Having established that H-CBASS2 provides defence against lytic infection, we next sought to assess whether it also mediates an antiviral response during chronic infection. To test this, we transformed the H-CBASS2 construct into *H. volcanii*, which also does not naturally encode any CBASS (Fig. 2A). We used the model virus HFPV-1, which causes a chronic, productive, non-lytic infection. HFPV-1 infects cells by attaching to the cell surface and delivering its genome into the host. The viral genome is then replicated without integration into the host genome. Newly assembled virions are subsequently released from the cell through a non-lytic, budding-like mechanism, allowing continuous virus production while the host cell largely remains alive [14]. We initiated experiments from virus-infected colonies (see methods Fig. 2A), examined the growth profile of the transformed cells and observed a very small degree of growth inhibition upon H-CBASS2 expression in *H. volcanii* compared to a negative control strain containing an empty vector, indicating a small fitness cost (Fig. 2B). Notably, virus-infected cells that were induced to express H-CBASS2 showed substantial growth inhibition compared to uninfected cells or to HFPV-1-infected cells carrying an empty vector (Fig. 2B, 2C and 2G). These results indicate that H-CBASS2 is active in *H. volcanii* and responds to chronic HFPV-1 infection.

**Figure 2.**
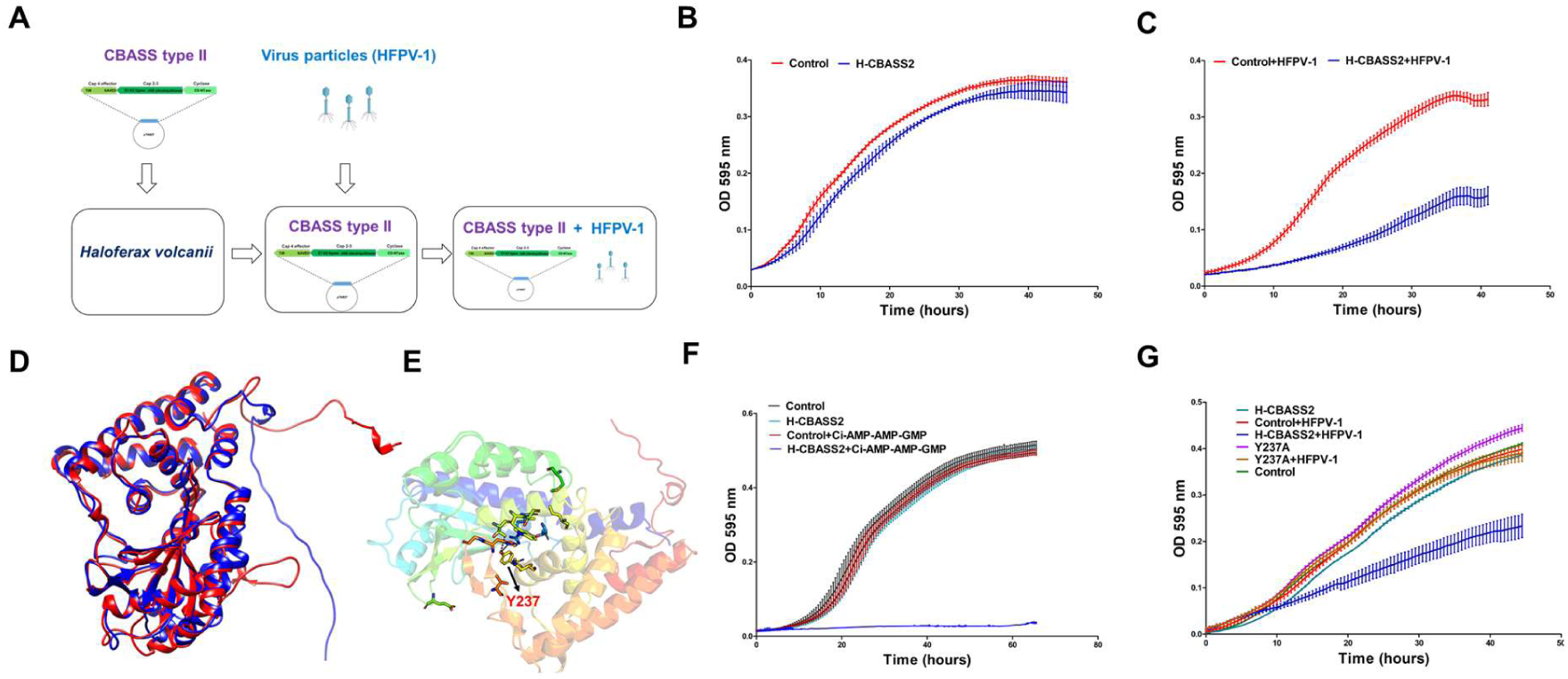
HFPV-1 infection causes growth delay in H-CBASS2 expressing cells. **A)** Outline of the HFPV-1 infection experiments. pCBASS2, a pTA927 plasmid encoding H-CBASS2 (see above) was transformed into *H. volcanii* (WR540). Subsequently, HFPV-1 particles were introduced into pCBASS2 containing *H. volcanii* cells, and virus-infected colonies were selected by PCR screening. **B)** Growth curves of *H. volcanii* expressing H-CBASS2 under tryptophan inducible promoter (blue) and control (with empty vector) (red), without viral infection. **C)** Growth curves of virus-infected H-CBASS2-expressing *H. volcanii* (blue) under tryptophan inducible promoter and infected control (red) are shown. **D)** 3D structural alignment of the predicted H-CBASS2 cyclase (AlphaFold 3) structure aligned with the crystal structure of the *E. cloacae* CBASS cyclase using Foldseek (TM score: 0.82595 and RMSD of 7.3 Å). Blue represents the H-CBASS2 cyclase and red represents the *E. cloacae* cyclase. **E)** The predicted 3D structure of the H-CBASS-2 cyclase is shown in cartoon representation, with residue Y237 highlighted in stick format (red) to indicate its position within the protein. **F)** Growth analysis of *H. volcanii* strains expressing H-CBASS2 (blue) and the empty vector control (red) in the presence of exogenous cyclic-AMP-AMP-GMP (12 μM). Untreated empty vector control and untreated H-CBASS2 strains are shown in black and cyan, respectively. **G)** Growth curves of mutated H-CBASS2 cyclase with or without virus infection. Y237A mutation on the H-CBASS2 cyclase without infection (purple) and with infection (brown), control in the absence (green) or the presence (red) of infection, wild-type H-CBASS2 without infection (cyan) and with infection (blue). B, C, F and G: Lines represent the mean of at least 3 biological replicates, each with three technical replicates.

### Characterization of H-CBASS2 signalling

A BLASTP analysis revealed that the H-CBASS2 cyclase had 42% sequence identity with the *Enterobacter cloacae* CBASS CDnD cyclase (Extended Fig. 1), which is known to produce cyclic AMP-AMP-GMP (cAAG) upon viral infection^17^. Additionally, we created a homology model of the H-CBASS2 cyclase using AlphaFold3, confirming that the aligned region has about 40% sequence identity and a TM-score of 0.8259 with the *E. cloacae* cyclase (Fig. 2D). Furthermore, when *H. volcanii* cells expressing either the full H-CBASS2 system or an empty vector control were treated with 6, and 12 μM of exogenous cAAG in the absence of viral infection, strong growth inhibition was observed specifically in H-CBASS2-expressing cells, with the most pronounced effect at 12 μM, compared to the empty vector control (Fig. 2F). In contrast, no growth inhibition was observed upon treatment with 12 μM of exogenous cyclic AMP-AMP-AMP (cAAA) and cyclic UMP-AMP (cUA) (Extended Fig. 2 a & b). This confirmed that the signal molecule that H-CBASS2 effector responds and is the most likely to be produced by its cyclase is cAAG.

Given the sequence, structure, and substrate similarity to the *E. cloacae* cyclase, we could predict the active site tyrosine at position 237 in the H-CBASS2 homolog and mutate it to alanine (Fig. 2E). This mutation was shown in the *E. cloacae* cyclase to completely abolish c-AAG synthesis, emphasizing the essential role of this residue in the cyclase activity ^17^. Indeed, the Y237A mutation in the H-CBASS2 cyclase rescued the substantial growth inhibition caused by HFPV-1 infection in the presence of H-CBASS2, as well as the minimal growth inhibition caused by H-CBASS2 alone (Fig. 2G). These results confirm that the H-CBASS2 cyclase, similar to *E. cloacae* CDnD, produces its signalling molecule in response to viral infection and is critical for activity.

### H-CBASS2 does not cause rapid cell death upon HFPV-1 infection

CBASS systems are considered to be abortive infection systems that kill the cell in order to stop viruses from spreading to neighbouring sister cells. Specifically, a bacterial type II CBASS system was shown to deplete cellular NAD^+^ levels and bring about cell death (or at least completely abolish growth) ^12^. We therefore performed live-dead staining assays on logarithmic cultures of HFPV-1-infected cells expressing H-CBASS2. Surprisingly, we detected very few dead cells, and their ratio in the culture did not exceed that observed in the vector-only control (Fig. 3A). Given that H-CBASS2 did not cause abortive infection in this case, we wondered whether it could nonetheless stop or delay the spread of HFPV-1 infection. HFPV-1 is a virus that does not lyse host cells and does not form plaques using standard approaches ^16^. However, since this virus does delay host growth, we used a recently developed protocol that enables the detection of plaques produced by non-lytic viruses of haloarchaea^18^. These plaques are caused by growth inhibition, as is commonly observed upon infection with non-lytic phages, such as M13 ^19^. Surprisingly, expression of H-CBASS2 did not substantially reduce the plating efficiency of HFPV-1 (Fig. 3B). Thus, H-CBASS2 probably cannot interfere with the early stages of infection by HFPV-1.

**Figure 3.**
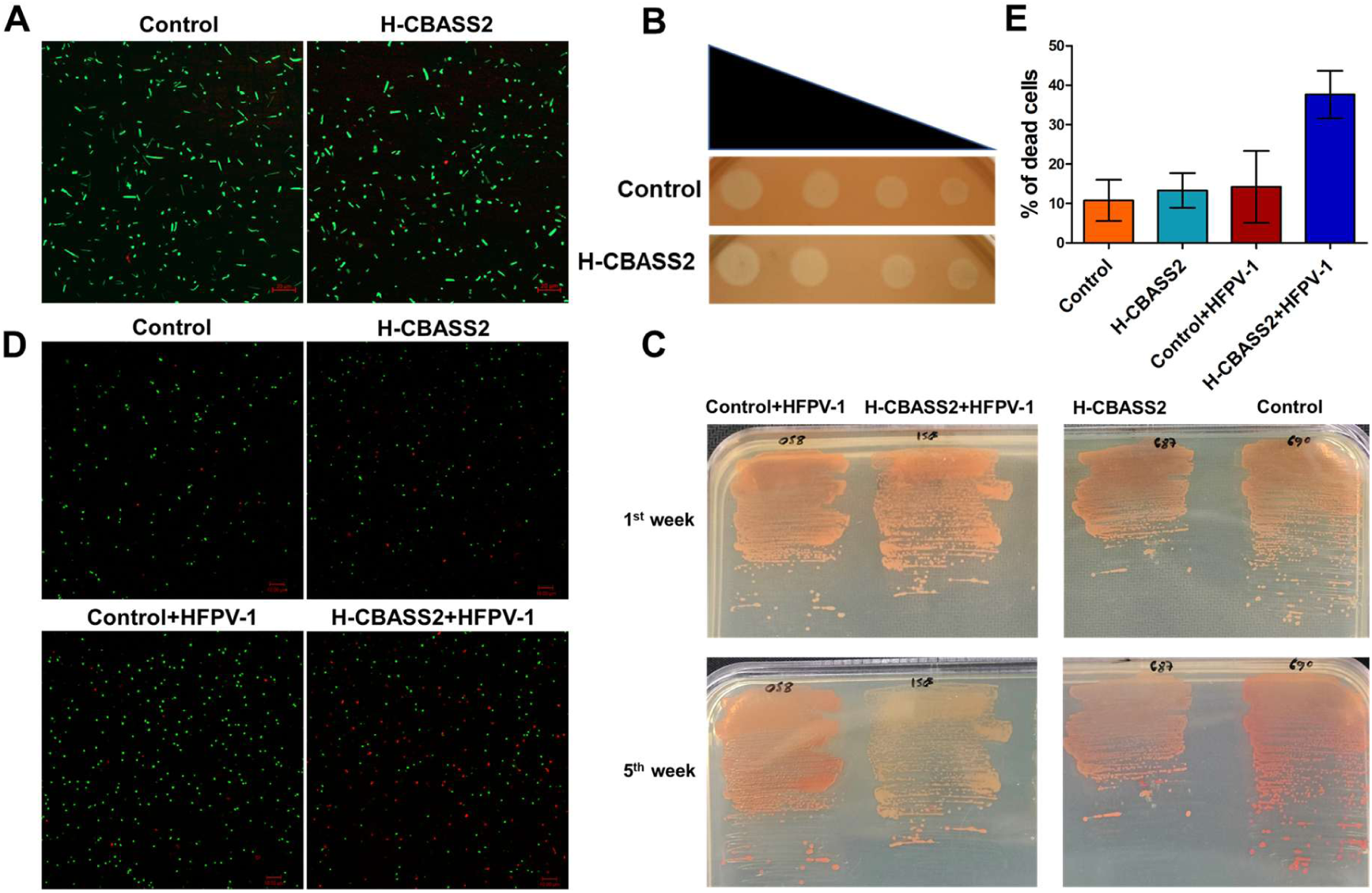
H-CBASS2 does not cause rapid cell death Upon HFPV-1 infection. **A)** Representative confocal microscopy images depicting Live/Dead assays of cells harvested during exponential growth (OD=∼0.6). H-CBASS2-expressing cells and control cells with an empty vector, both infected with HFPV-1 (Scale bar: 20 µm). Data are representative of at least 3 independent experiments. **B)** Plaque assay using 10-fold serial dilutions of virus (left-most is a 10^−4^ virus dilution suspension). H-CBASS2 expressing cells and control cells with an empty vector were mixed with 0.02% top agar and plated onto rich medium. Afterward, 3 µL of serial dilutions of HFPV-1 were spotted to quantify plaquing efficiency. The images are representative of at least three biological replicates. **C)** Representative images of colony colour of *H. volcanii* cells infected with HFPV-1. colonies were streaked on casamino agar for *H. volcanii* strains expressing H-CBASS-2, with and without virus infection, along with a control containing an empty vector. A 20 µl exponentially growing culture was streaked onto fresh plates, which were incubated at 45°C for 4-5 days. After growth became visible, the plates were kept at room temperature for 4-6 weeks, with images taken every 3 days. Virus-infected H-CBASS2-expressing colonies left at room temperature for 4-6 weeks showed noticeable bleaching. The images are representative of at least three biological replicates **D)** Live/Dead assays conducted on cells from bleached colonies, revealing a higher number of dead cells in the virus-infected *H. volcanii* strain with H-CBASS-2 compared to the control (Scale bar: 10 µm). The images are representative of at least three biological replicates. **E)** Quantification of Live/Dead assays from bleached colonies (N = 2). In both Figures 3A and 3D, green fluorescence represents live cells and red fluorescence represents dead cells.

### Long-term exposure to viral infection in HCBASS-2-expressing cells reduces viability

Although we did not detect any short-term cell mortality, we did observe that virus-infected H-CBASS2-expressing colonies left at room temperature for 4-5 weeks became bleached (Fig. 3C, see also time series in Extended Fig. 3), losing the pink-red pigmentation typical of Haloferax cells that naturally synthesize the carotenoid pigment bacterioruberin^20,21^. When suspending cells from the bleached colonies and submitting them to Live-Dead assays, we observed that many of the cells from those colonies were dead. In contrast, 4–5-week-old colonies of infected cells that did not express H-CBASS2 maintained normal colour and viability (Fig. 3D & 3E). Importantly, carotenoids as well as essential archaeal membrane lipids both share a key building block, mevalonate, synthesized from HMG-CoA by the enzyme HMG-CoA reductase that uses two molecules of NADPH^22^. Thus, it is possible that the depletion of cellular NAD (and NADPH) will force cells to scavenge mevalonate from carotenoids into the more critical membrane lipids, resulting in the observed bleached phenotype.

### HCBASS-2-expressing cells require long-term exposure to HFPV-1 to deplete cellular NAD^+^

Given the NAD depletion observed during HFTV1 infection of H-CBASS2-expressing cells, we also quantified total NAD levels during HFPV-1 infection of cells harbouring H-CBASS2 (see Methods). Surprisingly, total NAD levels during HFPV1 infection were largely unaffected in infected H-CBASS2-expressing cells during logarithmic growth compared to controls with an empty vector (Fig. 4A). In contrast, during late logarithmic phase, we observed a slight reduction in NAD levels in the H-CBASS2-expressing cells (Fig. 4B). Furthermore, when we measured NAD levels from the bleached colonies described above in figure 3C, we observed that NAD levels were greatly decreased in virus-infected *H. volcanii* with H-CBASS2 (Fig. 4C) compared to infected cells with the empty vector, comparable to those observed during HFTV1 infection (Fig. 1E). We also observed a modest decrease in NAD levels in H-CBASS2-expressing cells from 4-week-old colonies that were not infected. These results suggest that NAD depletion in *H. volcanii* cells by H-CBASS2 upon chronic HFPV-1 infection is a slow process and becomes lethal only after many generations of growth.

**Figure 4.**
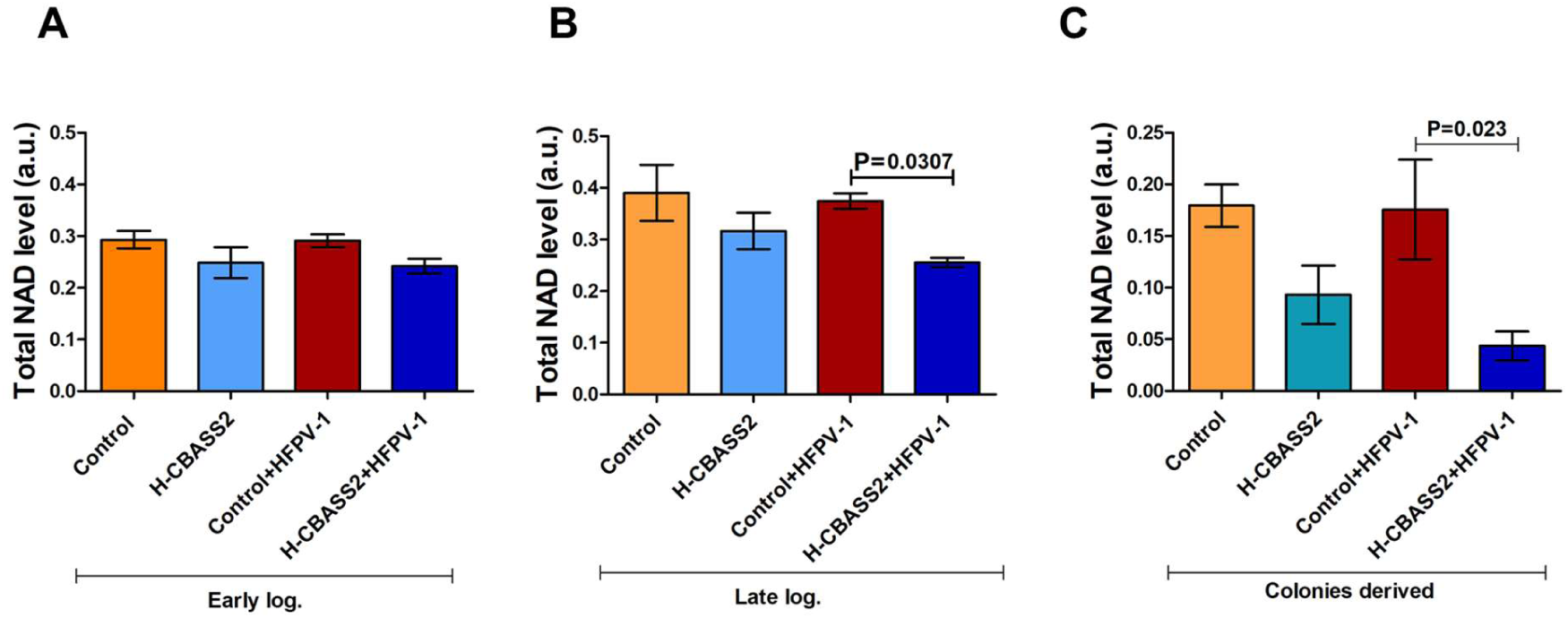
NAD levels in HCBASS-2-expressing cells only decrease after sustained virus exposure. **A)** NAD levels in *H. volcanii* strains expressing H-CBASS-2, in the presence or absence of virus infection. Strains were grown to early exponential growth phases. **B)** NAD levels in *H. volcanii* strains expressing H-CBASS-2, in the presence or absence of virus infection. Strains were grown to late exponential growth phases. Equal numbers of cells, normalized by OD, from each culture were collected and total NAD levels were measured using the NAD/NADH Quantification Kit (panels A and B). Data are representative of at least three independent experiments. NAD⁺ levels in Figure 4B were compared between strains using a paired-sample t-test. Control + HFPV-1 vs. H-CBASS2 + HFPV-1: *P* = 0.0307; CBASS2 vs. H-CBASS2 + HFPV-1: *P* = 0.174. **C)** NAD levels in cells from bleached colonies. After resuspending the bleached cells (previously incubated for 4-5 weeks in room temperature) in fresh medium, protein concentration of the biomass was determined using the Bradford method. A biomass with equal protein concentration from each strain was then used to measure total NAD levels. Data represent results from at least three independent experiments. NAD⁺ values were compared between strains using a paired-sample t-test. H-CBASS2 vs. H-CBASS2+HFPV-1: P = 0.080; Control+HFPV-1 vs. H-CBASS2+HFPV-1: P = 0.023; Control vs. H-CBASS2: P = 0.111.

### H-CBASS2 eradicates HFPV-1 infection

Since HFPV-1 infection is relatively mild while H-CBASS2 activity can be harmful and even lethal, the question arises whether such a system benefits the host during chronic infection, or whether it is deleterious. Accordingly, we performed a head-to-head competition experiment in which we grew together HFPV-1-infected *H. volcanii* cells, with or without H-CBASS2. As expected, at 24 and 48 hours after mixing, the H-CBASS2-negative cells increased in relative abundance at the expense of the H-CBASS2-positive cells (Fig. 5A), indicating that in the short-term H-CBASS2 is detrimental during chronic infection in an unstructured environment. However, in their natural environment, Haloferax species often create structured communities, such as biofilms, where all cells in a biofilm patch are sister cell derived from the same clone, and therefore *either* CBASS-positive or CBASS negative. We thus tested what occurs during prolonged chronic infection of H-CBASS2-expressing cells. We performed an in-vitro evolution experiment where we grew HFPV-1-infected *H. volcanii* cells with and without H-CBASS2 separately in liquid culture until the late stationary phase, and then used that culture to inoculate fresh medium and repeated this for ∼10 passages, taking samples at passages 3, 6, and 10 (Fig. 5B). We then plated the diluted cultures from these time points on solid media and screened 30-40 colonies for the presence of HFPV-1 (Extended Fig. 4a and 4b for PCR screen gels). Remarkably, nearly all colonies from H-CBASS2-expressing cells no longer had detectable virus DNA at passage 10, in all three biological replicates, while over 95% of colonies that did not express H-CBASS2 still had viral DNA presence (Fig. 5C, D, and E). Even as early as passage 3 (less than 15 generations), most H-CBASS2-expressing colonies did not have detectable viral DNA Extended Fig. 4d and e for PCR screen gels & Fig. 5C). In comparison, the presence of the H-CBASS2 insert was confirmed by PCR after the 10th passage in all colonies from the H-CBASS2-expressing culture (Extended Fig. 4c). Sequencing of several colonies after passage 10^th^ did not detect any mutation in the H-CBASS2 or the plasmid vector that harbours it.

**Figure 5.**
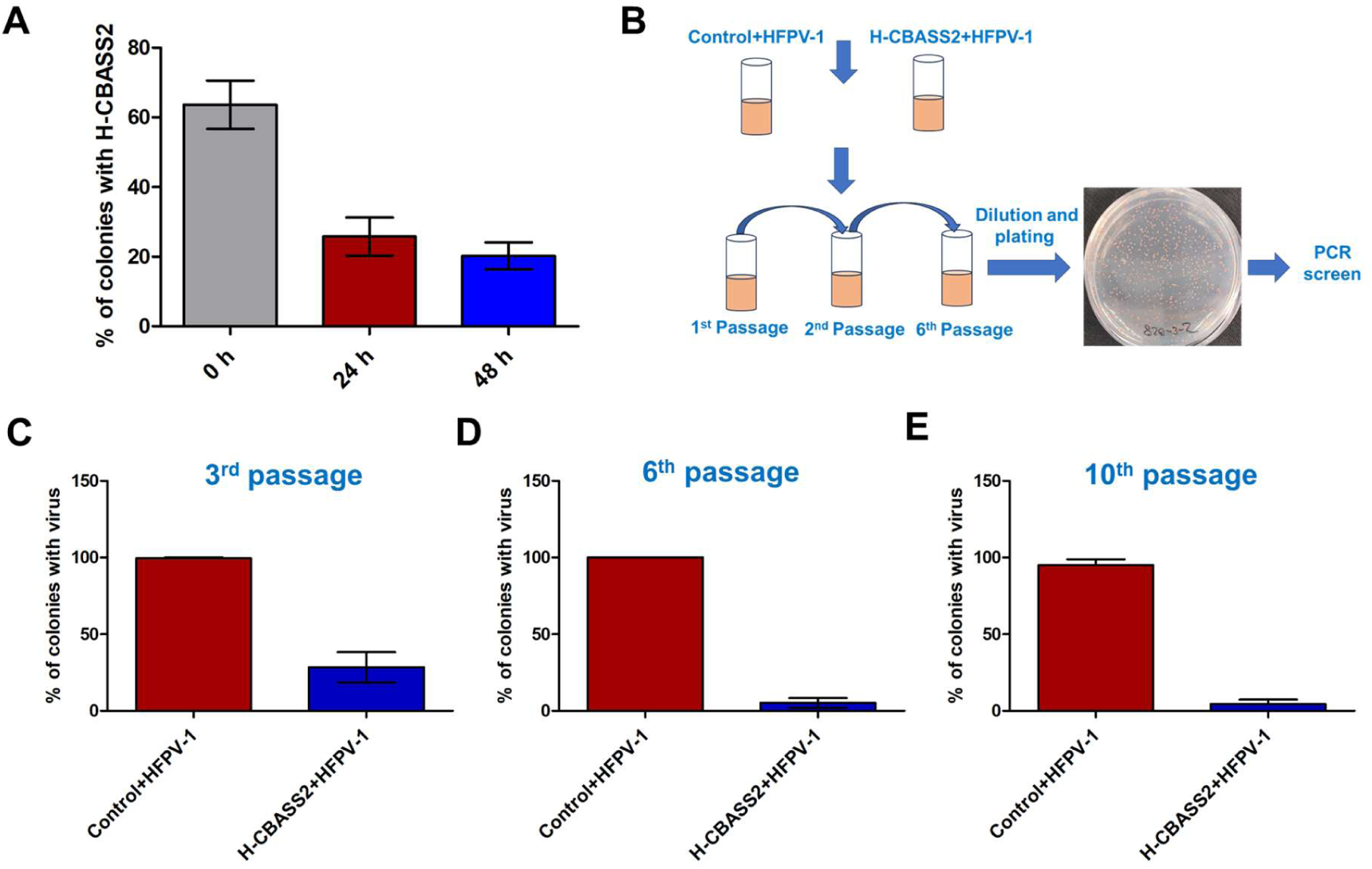
H-CBASS2 clears HFPV-1 infection. **A)** Head-to-head competition experiments between a virus-infected *H. volcanii* strain carrying the H-CBASS-2 system and a virus-infected control strain lacking that system. Equal amounts of log-phase cultures were mixed into fresh Hv-ECas medium and incubated at 45°C for 48 hours. Samples of 100 µL were taken at 24 and 48 hours and plated on Hv-ECas plates. Once colonies appeared, the presence of CBASS was tested by PCR. Each strain was tested with at least 3 biological replicates. **B)** Schematic representation of an in vitro evolution experiment. In this experiment, a virus-infected *H. volcanii* strain expressing H-CBASS-2 and a control strain infected with the virus were used. Single isolated colonies from each strain (infected with HFPV-1) were inoculated into fresh Hv-ECas medium supplemented with thymidine and tryptophan. The cultures were then serially passaged approximately 10 times in fresh medium. Afterward, the cells were plated on Hv-ECas medium supplemented with thymidine and tryptophan, and the presence of virus DNA was tested by PCR. Three biological replicates were performed. **C)** Ratio of virus-cured colonies after the 3^rd^ passage. **D)** Ratio of virus-cured colonies after the 6^th^ passage. **E)** Ratio of virus-cured colonies after the 10^th^ passage.

In addition, the virus-cured H-CBASS2–expressing strains obtained after the 10th passage exhibited a slight growth advantage compared with both control cells carrying an empty vector and H-CBASS2–expressing that were never exposed to viruses (Extended Fig. 5). In contrast, during infection (prior to viral clearance), H-CBASS2-expressing cells exhibited markedly reduced growth (Fig. 2C & 2G). This implies that most of the cells cleared the virus and probably acquired mutations during infection (see below).

We then performed qPCR to quantify the level of virus DNA in the supernatant after 48 and 72 hours of growth and observed almost a 20% decrease in viral DNA in the supernatant in the CBASS-expressing cells compared to the control infected cells. Moreover, we also observed 20% lower levels of viral DNA after the 4th passage and 40% lower levels of viral DNA after the 6th passage in the CBASS-expressing cells, which can probably be attributed to virus clearing in the majority of cells at these later stages (Extended Fig. 6a, b, c, and d). Taken together these findings indicate that viral production declines and viruses are gradually eliminated in the CBASS-expressing cultures.

### H-CBASS2 expression disrupts a key host protein and alters viral gene expression

Next, we wanted to explore how H-CBASS2-expressing cells eliminate viruses. Recent studies have demonstrated that some defense systems can modify viral proteins and interfere with phage assembly ^23^ and egress ^24^. To determine whether H-CBASS2 affects viral egress, we purified HFPV-1 from the supernatant of both H-CBASS2–expressing and control cells after 7 days of continuous growth and examined viral particles by TEM. No differences in virion morphology were detected in H-CBASS2–expressing cells. In addition, plaque assays performed with these viruses on the same *H. volcanii* UG690 reporter strain showed that viruses derived from H-CBASS2–expressing cells formed plaques with efficiencies comparable to those of the non CBASS-exposed controls (Extended Fig. 7). Taken together, these results suggest that H-CBASS2 does not directly damage viral particles.

We then performed proteomic analysis on HFPV-1-infected cells, both with and without H-CBASS2 expression from stationary phase cultures. Of the eleven predicted viral proteins, we identified six, with several showing substantial differences in abundance between H-CBASS2-expressing and control cells strains (Table 1). Most notably, a putative transcriptional regulator (UVW56703.1) was undetectable in H-CBASS2–expressing cells, whereas the viral spike protein was more than two-fold higher. We then repeated the experiment, this time using late–log-phase cultures. The same proteomic pattern for HFPV-1 was observed in these cultures, including complete loss of UVW56703.1 in all H-CBASS2–expressing cells (Extended Table 1). Furthermore, when we tested viral gene transcription using qRT-PCR and observed an approximately 2-fold decrease in UVW56703.1 transcript in log-phase H-CBASS2-expressing cells (Extended Fig. 8). Taken together, these findings strongly suggest that H-CBASS2 limits HFPV-1 infection by disrupting expression of specific viral proteins rather than by blocking particle assembly or egress.

**Table 1.**
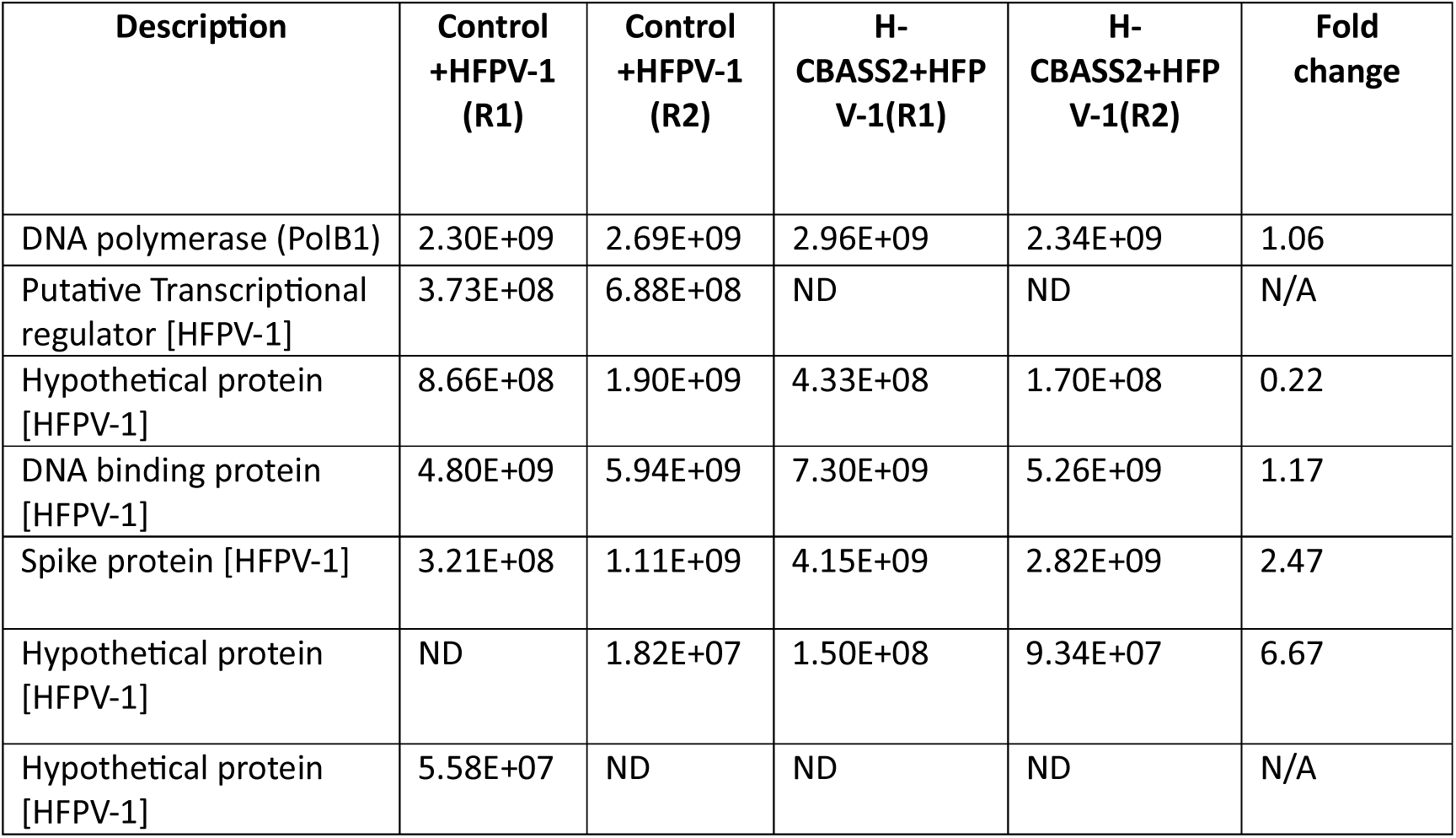
Abundance of HFPV-1 proteins during H-CBASS2 expression. HFPV-1 proteins detected in infected *H. volcanii* cells by MS/MS. The “Fold Change” column represents the ratio of the mean of two biological replicates (**R1 - Replicate 1, R2 - Replicate 2**) of infected control and H-CBASS2 cells. The host housekeeping gene **DNA polymerase B (*polB1*)** is shown as a control since its expression is expected to be unaffected by H-CBASS2.

| Description | Control +HFPV-1 (R1) | Control +HFPV-1 (R2) | H-CBASS2+HFP V-1(R1) | H-CBASS2+HFP V-1(R2) | Fold change |
| --- | --- | --- | --- | --- | --- |
| DNA polymerase (PolB1) | 2.30E+09 | 2.69E+09 | 2.96E+09 | 2.34E+09 | 1.06 |
| Putative Transcriptional regulator [HFPV-1] | 3.73E+08 | 6.88E+08 | ND | ND | N/A |
| Hypothetical protein [HFPV-1] | 8.66E+08 | 1.90E+09 | 4.33E+08 | 1.70E+08 | 0.22 |
| DNA binding protein [HFPV-1] | 4.80E+09 | 5.94E+09 | 7.30E+09 | 5.26E+09 | 1.17 |
| Spike protein [HFPV-1] | 3.21E+08 | 1.11E+09 | 4.15E+09 | 2.82E+09 | 2.47 |
| Hypothetical protein [HFPV-1] | ND | 1.82E+07 | 1.50E+08 | 9.34E+07 | 6.67 |
| Hypothetical protein [HFPV-1] | 5.58E+07 | ND | ND | ND | N/A |

We also analyzed host protein abundance in virus-infected cells with or without H-CBASS2 expression. We observed that several host proteins involved in carbohydrate metabolism, transport, and host signaling and regulation were markedly less abundant in virus-infected cells expressing H-CBASS2 compare to infected control. Notably, these proteins were largely unaffected when comparing uninfected cells with or without H-CBASS2 expression. (Extended Fig. 9) This suggests that H-CBASS2 expression primarily impacts host metabolism of virus-infected cells. A likely explanation is that upon infection, H-CBASS2 activity, primarily NAD^+^ depletion, alters cellular metabolism, thereby limiting the cell’s capacity to support viral protein synthesis. As a result, viral replication may be progressively impaired rather than abruptly blocked, which could explain why we do not observe a strong direct effect on virus particle production or a pronounced difference in plaquing efficiency. Collectively, these findings support a model in which H-CBASS2 exerts its antiviral effect indirectly by disrupting the cellular metabolic environment rather than by direct modification of viral proteins.

### H-CBASS2-containing strains that cleared HFPV-1 infection exhibit resistance to re-infection by HFPV-1

An additional mechanism that can lead to virus eradication is host evolution of resistance towards that virus. We hypothesized that cells that eliminated the virus might develop resistance and eventually outcompete the virus-sensitive population. Cells that spontaneously clear the virus will grow much faster than infected ones, since they do not suffer from the effects of H-CBASS2 on growth, but unless they have virus-resistance mutations they will quickly be re-infected. To test this hypothesis, we selected four independent colonies that encode H-CBASS2 and had been cured of the virus (after the 10^th^ passage), and performed a plaque assay on colony-derived cells. Indeed, all four virus-cured strains exhibited lower plating efficiency compared to strains derived from the control and H-CBASS2-positive colonies that had never been infected with the virus (Fig. 6A and B). To test whether this resistance emergence was specific, we passaged both the strain expressing H-CBASS2 (without viral infection) and wild-type control cells with an empty vector through ten passages. We then selected two to three colonies from each condition and performed plaque assays. The results showed no difference in plaque formation between uninfected H-CBASS2-expressing cells before and after passaging, and the same was observed for the empty vector controls. We therefore conclude that prior HFPV-1 exposure enables *H. volcanii* cells carrying H-CBASS2 to acquire stable, heritable HFPV-1 resistance. Curiously, we also observed that these virus-cured strains exhibited a markedly different colony appearance, characterized by darker pigmentation, smaller size, and a smoother, slimier surface then w.t. *H. volcanii* (Fig. 6C), leading us to speculate that these mutants have modified surface properties, as is often the case for phage-resistant surface mutants as well as virus-resistant mutants in archaea^25^. When we exposed these mutants to the H-CBASS2 signal molecule cAAG we observed a resistance phenotype, which varied among them (Extended Fig. 10). This phenotype could indicate resistance to CBASS activity, but could also reflect reduced uptake of the signal due to altered envelope properties.

**Figure 6.**
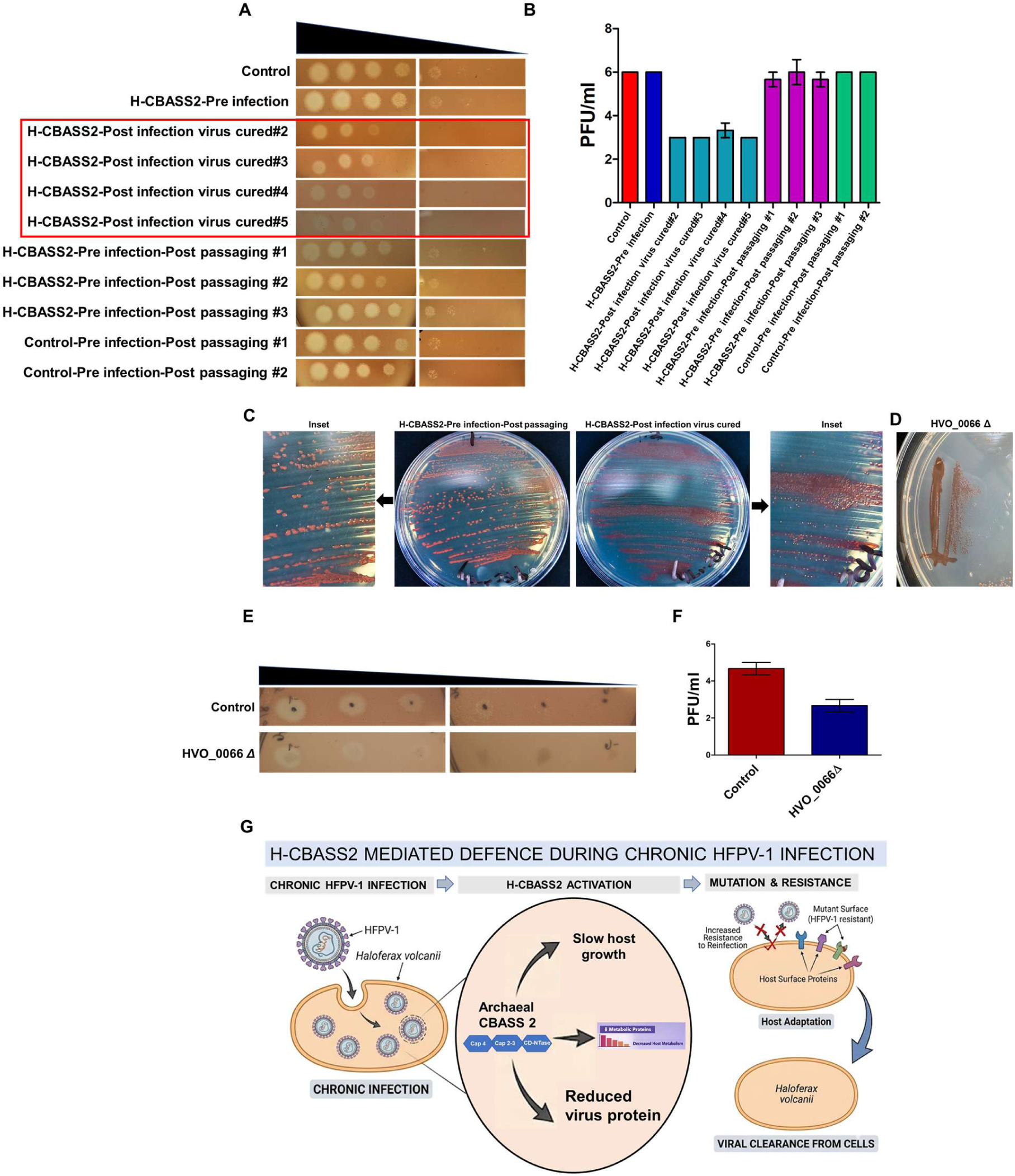
Virus-cured Haloferax strains expressing H-CBASS2 show resistance to re-infection by HFPV-1. **A)** Plating efficiency of HFPV-1 on different *H. volcanii* strains: H-CBASS2-expressing post-infection-post-passaging virus-cured colonies (marked in red), together with the pre-infection (never-infected) H-CBASS2-expressing strain and the non-infected *H. volcanii* strain carrying an empty vector as controls. Also shown are uninfected H-CBASS2-expressing cells and the empty-vector control after passaging. Ten-fold serial dilutions of HFPV-1 were spotted onto the plates (left-most is undiluted). Each strain was tested with at least 3 biological replicates. **B)** Quantification of plaque assays of Figure 6 A (N = 3).**C)** Representative images of colony morphology across different strains. H-CBASS2 post-infection after passaging virus-cured strains exhibit distinctly smaller colonies and darker pigmentation compared to H-CBASS2 pre-infection-post-passaging strains and wild-type *H. volcanii*. **D)** Representative images of colony morphology of HVO_0066 transposon mutant. **E)** Images represent the plating efficiency of HFPV-1 on HVO_0066 disruption strains. Ten-fold serial dilutions of HFPV-1 were spotted onto plates, with the leftmost spot representing the undiluted virus sample. Each strain was tested in at least three biological replicates. **F).** Quantification of plating assay of Figure 6E (N = 3). **G)** Proposed mechanism of CBASS2-mediated activity against HFPV-1 infection. Chronic infection of *H. volcanii* by HFPV-1 triggers activation of the archaeal CBASS2 system, initiating a multi-pronged defensive response. This signalling leads to gradual NAD^+^ depletion followed by a marked decrease in host metabolic proteins, which slows host cell growth and concurrently reduces intracellular viral protein levels. Over time, host adaptation occurs through mutations in surface proteins, increasing resistance to reinfection. The synergy between suppressed viral replication and prevention of re-entry ultimately results in complete clearance of HFPV-1 from the cell population.

### Lipoprotein mutations confer resistance to HFPV-1

To identify genomic events that could explain this resistance, we extracted DNA from both H-CBASS2-expressing virus-cured colonies and virus-infected control colonies. We then sequenced the genomes from four colonies of each type obtained at passage 10 in order to infer what could have mediated viral clearance. Previous studies have shown that some defense systems, which induce dormancy rather than cell death, allow CRISPR-Cas sufficient time to acquire spacers from the viral invader’s genome, and subsequently, CRISPR-Cas degrades the viral genome. *H. volcanii* has an active CRISPR-Cas system ^26–28^, yet past work has shown CRISPR-Cas to be unable to clear HFPV-1 infection^16^. Analysis of the CRISPR arrays of the H-CBASS2-expressing clones that cleared the infection showed that no new spacers were acquired, indicating that HFPV-1 eradication probably did not involve CRISPR-Cas activity. Additionally, no mutations in the plasmid from which H-CBASS2 was expressed were detected, nor was any HFPV-1 virus presence were detected in the cured strains.

Next, we identified several point mutations in the genomes of the virus-resistant colonies, some of which occurred in two or more clones derived from different colonies (Extended Table 2). Notably, point mutations in loci associated with lipoproteins were observed in nearly all virus-cured strains. For example, a single nucleotide deletion - located in the non-coding region upstream of a putative lipoprotein-associated gene (HVO_2285) in *H. volcanii* was shared among 3 out of 4 virus-cured strains. However, mutations were also detected in the same non-coding region in two CBASS-negative samples, though at different positions. Interestingly, this non-coding region is located within a putative endogenous provirus of *H. volcanii* region known as Halfvol4^29^ (Extended Table 2)^30^. In addition, we identified a single synonymous substitution in HVO_2266 an uncharacterized protein also located within a putative endogenous provirus in two control samples. Thus, mutations in this locus probably occur during HFPV-1 infection in both CBASS-positive and CBASS-negative cells.

Furthermore, we detected a frameshift mutation in HVO_2141, which encodes a putative lipoprotein, in two independent virus-cured strains. We also identified a single nucleotide substitution in two virus-cured strains in the putative lipoprotein HVO_0066, which contains several PQQ repeat domains, a lipobox motif, and a signal peptide sequence typical to for twin-arginine transporter secretion (TAT)^31,32^ (Extended Table 2).

Notably, none of the latter mutations were observed when we sequenced uninfected H-CBASS2-expressing cells or wild-type control cells with an empty vector after ten passages. Furthermore, no similar or specific mutations were identified in H-CBASS2-expressing colonies prior to passaging (Extended Table 2, Sample-9-10). Mutations affecting lipoproteins might alter the surface and disrupt viral adherence or entry, which could explain the observed resistance to reinfection. However, all these mutants had multiple mutations in their genome, making it difficult to ascertain this interpretation of the findings. We therefore obtained a transposon insertion mutant in the HVO_0066 gene in wild-type *H. volcanii* DS2 that was never exposed to HFPV-1. Interestingly, this strain displayed colonies with darker pigmentation and smaller size, similar to the virus-cured mutant strains (Fig. 6D). Plaque assays with HFPV-1 confirmed that this strain was as resistant to HFPV-1 infection as the CBASS-expressing cells that eliminated the virus after passaging (Fig. 6E & F). Altogether, these results suggest that this lipoprotein likely serves as a viral receptor or is required for successful viral entry, and its loss results in concomitant changes to both viral susceptibility and colony morphology. Thus, H-CBASS2 appears to function as a two-stage defence in this case: disrupting viral gene expression, while simultaneously creating a burden on growth that provides a large advantage to cells that cleared the virus. Such cells will be quickly re-infected by neighbours, unless they have also acquired resistance-conferring mutations in surface-associated genes. Such resistant mutants that are only rarely infected can then quickly take over the culture by outcompeting w.t. HFPV-1-infected sister cells that are also burdened by CBASS activity.

## Conclusion

Here, we characterize the Type 2 CBASS system from the halophilic archaeon *Haloferax* strain Atlit 48N in two archaeal virus-host models. Intriguingly, although HFPV-1 and HFTV1 do not have any protein in common, and their mode of infection is completely different, H-CBASS2, nonetheless responded to both of them, which begs the question - what does the system sense? Two explanations come to mind: either H-CBASS2 senses a replication intermediate that is shared between these viruses (such as the 3′ ssDNA ends sensed by Hachiman type I-A systems)^33^, or it responds to a host response molecule that is markedly increased in both infections. Establishing which of these mechanisms, if any, underlies H-CBASS2 activity will require further study.

In the HFTV1-*H. gibbonsii* infection model, H-CBASS2 caused pre-mature lysis of cells, thereby reducing the efficiency of infection. It is likely that NAD^+^ depletion aggravates membrane stress caused by HFTV1 infection because NADPH is so critical in archaea for biosynthesis of membrane lipids. In contrast, because HFPV-1 is non lytic this does not occur in the *H. volcanii*-HFPV-1 system, where activity is also much slower (see total NAD levels in figure 4 A).

Although we were skeptical about H-CBASS2 being beneficial under chronic virus infection, we nevertheless observed that H-CBASS2-expressing cells cleared HFPV-1 infection: this is probably a combination of a direct effect on HFPV-1, and reduced toxicity to the host, which increases the growth advantage of cells that eliminate the virus (which may also be more tolerant to CBASS activity itself. When such cells are also genetically resistant to re-infection, they will inevitably dominate and drive the virus extinct. Our findings highlight a multifactorial mechanism of virus eradication involving both defense system activity and mutation-based adaptation. Our results support the broader view of so-called abortive-infection systems as defenses that target the virus as well as the host, and can affect infection outcome long before causing cell death^34^. Indeed, a recent study in the bacterium *Pseudomonas aeruginosa* showed a phospholipase-based CBASS capable of protecting against multiple phages with no detrimental effect on cell growth^35^, representing an even more extreme example than ours. We therefore conclude that CBASS function can be anywhere on a spectrum that spans abortive infection on one extreme and completely virus-specific damage on the other, depending on the virus, the system (and its expression level)^36^, and the host.

## Methods

### Culture conditions

The Haloferax wild-type strains were routinely cultured at 45°C, either in Hv-YPC or Hv-Cas/Hv-Enhanced Cas medium. *Haloferax volcanii* transformants were selected for and grown either in Hv-Cas or Hv-Enhanced Cas (Hv-ECas). Thymidine 40 μg/ml and tryptophan 50 μg/ml were supplemented when required. Bacterial strains, were cultured at 37°C in LB medium or LB medium supplemented with ampicillin for strains carrying plasmids.

### Cloning and mutagenesis

Complete *Haloferax* 48N CBASS Type II system, was cloned into high-copy replicating plasmids pTA927 (which contains the tryptophanase inducible promoter) by using the Gibson assembly method. DNA fragments for the DNA inserts, such as H-CBASS2, and the plasmid vectors were first PCR-amplified with specific primers by using either Phusion or KAPA-HiFi DNA-Polymerase. PCR amplified plasmids and DNA fragments were then purified by the Wizard® SV Gel and PCR Clean-Up kit (Promega) followed by *Dpn*I digestion of purified plasmids. The purified DNA fragments and plasmids were then ligated using the Gibson assembly protocol^37^ (Gibson et al., 2009). After ligation, the resulting plasmids were transformed into *E. coli* DH12S cells using the electroporation method. Following the clone confirmation by PCR, the plasmids were extracted using the GenElute™ Plasmid Miniprep Kit from Sigma-Aldrich and transformed into *H. volcanii* strains.

### Mutagenesis

The Cyclase-dead variant of H-CBASS2 cloned into pTA927 was generated by substituting the conserved tyrosine amino acid at position 237 with alanine using site-directed mutagenesis, and the mutation was verified through sequencing.

### Transposon insertion mutant

The transposon (Tn) mutant used in this study is part of a genome-wide insertion library generated in *H. volcanii* as previously described^38^. Briefly, the library was constructed using an in vitro transposition approach with a mariner-based transposon system to generate random insertional disruptions across the genome. The HVO_0066 Tn insertion mutant was identified in a hypermotility screen of this library, and whole-genome sequencing revealed that the transposon is inserted at nucleotide position 267 of that gene.

### Transformation with HFPV-1

*H. volcanii* strains were grown overnight to the log phase and then harvested by centrifugation at 6,500 rpm for 5 minutes. The pellet was gently resuspended in 200 µl of spheroplasting solution containing 1 M NaCl, 27 mM KCl, 50 mM Tris-HCl, and 15% sucrose. The solution was treated with 0.5 M ethylenediaminetetraacetic acid (EDTA) at pH 8 to chelate any divalent cations present. After incubating for 10 minutes at room temperature, 5-10 µl of concentrated HFPV-1 particles were added to the resuspended culture, followed by another 5-minute incubation at room temperature. After the virus particles were added, 250 μL of 60% PEG600 was gently mixed into the solution. The mixture was then incubated at room temperature for 1 hour. Following the incubation with PEG600, the cells were washed with 1 ml of regeneration solution (Hv-YPC+ media with 15% sucrose). The pellets were then dissolved in Hv-YPC+ media with 15% sucrose and incubated at 28°C for 3 hours without shaking. After the 3-hour incubation at 28°C, the cells were transferred to a 28°C incubator shaker for an additional 3 hours. Subsequently, the cells were serially diluted and plated to obtain single colonies. HFPV-1-infected colonies were then identified by PCR using virus-specific primers.

### Growth curves

For all the growth experiments, each strain was grown over-night to the log phase in Hv-ECas medium with thymidine and tryptophan and then diluted into a fresh Hv-ECas medium with thymidine and tryptophan to OD0.03 to 0.05 and further kept at 42°C for 72 h with continuous shaking in 96-well plates. For induction, L-tryptophan (Sigma Aldrich) was added to the diluted culture to achieve a final concentration of 2 mM. Turbidity of the culture (OD595nm) was measured every 30 minutes using a microplate reader (Biotek ELX808IU-PC). For each strain, we performed a minimum of three biological replicates, with each biological replicate comprising three technical replicates. **Growth curves with HFTV1 —** *H. gibbonsii* strains expressing H-CBASS2 under a tryptophan-inducible promoter and the empty vector control were grown overnight to logarithmic phase in Hv-ECas medium. Cultures were then diluted into fresh Hv-ECas medium supplemented with 2 mM tryptophan to an OD of 0.05, after which HFTV-1 was added at an MOI of 0.01. Cultures were then incubated at 42°C for 72 hours with continuous shaking in 96-well plates. **Growth curve analysis with exogenous cyclic signaling molecules** To assess the effect of exogenous cyclic signaling molecules on H-CBASS2 activity, growth experiments were performed using uninfected H*. volcanii* cells expressing H-CBASS2 and an empty vector control strain. Both strains were grown overnight in Hv-ECas medium supplemented with thymidine and tryptophan until reaching log phase, then diluted to an OD₆₀₀ of 0.05 in fresh medium containing the indicated concentrations (6–12 μM) of cyclic molecules, including cyclic-AMP-AMP-GMP (cAAG), cyclic-AMP-AMP-AMP (cAAA), and cyclic-UMP-AMP (cUA). Cultures were incubated at 45°C with continuous shaking for 72 hours, and growth was monitored by measuring optical density at regular intervals as described above.

### Live/Dead staining

To evaluate the viability of virus-infected *H. volcanii* strains expressing H-CBASS-2, we conducted a Live/Dead staining experiment using the LIVE/DEAD BacLight Bacterial Viability Kit from Thermo-Fisher (Ref-L13152). To determine the ratio of dead cells from the logarithmic phase culture, each strain was grown overnight. The cultures were then diluted into fresh medium to an optical density (OD) of 0.03 to 0.05 and incubated in a 45°C shaking incubator until they reached the logarithmic phase. Following that, 100 µl of the resuspended cultures were aliquoted into 1 ml Eppendorf tubes for subsequent analysis. Next, 10 μl of Propidium iodide dye and 10 μl of SYTO9 dye were added separately from their respective stock solutions. The treated cultures were then incubated in the dark for 15-20 minutes to allow for the cellular uptake of the stains. After the incubation period, 10 μl of the treated culture were placed onto a glass slide and examined using a confocal microscope (Leica sp8 confocal microscope) at magnifications of 20x and 40x. To evaluate dead cells from the bleached colonies, cells were directly taken from the plates showing bleached colonies and mixed into 100 μl of Hv-ECas medium. Subsequently, the mixed cells were washed twice with Hv-ECas medium and then subjected to the aforementioned microscopy protocol.

### Live/Dead assay of *H. gibbonsii* during HFTV1 infection

Overnight starter cultures of the empty vector control strain (UG788) and the H-CBASS2-expressing strain (UG784) were inoculated into Hv-ECas medium and grown in triplicate at 45°C with shaking. Upon reaching an OD₆₀₀ of 0.8–0.9, cultures were diluted to an OD₆₀₀ of 0.4 in 2 ml of fresh ECas medium supplemented with tryptophan. For each strain, two tubes were prepared: one for virus infection and one as an uninfected control. Infection was initiated by inoculating the appropriate tubes with HFTV1 at a MOI of 0.05. All tubes were subsequently incubated at 37°C for an additional 3–6 hours, after which the live/dead staining protocol was performed as described above.

### Plate colony pigmentation assays

For these plate assays, each strain was grown overnight to the log phase and then diluted to a fresh medium to OD 0.05, 20 µl of sample streaked on the new plates and kept at 45°C for 4-5 days. After the cells grew, all the plates were left at room temperature for 4-6 weeks, and images were captured every three days.

### NADH/NAD assay

Total cellular NAD level was measured using the NAD/NADH Quantification Kit from Sigma-Aldrich (MAK037). Each strain was grown to the late log phase, and then a roughly equal number of cells was taken from each culture, estimated by optical density. The cells were washed twice with cold Hv-Cas media. The pelleted cells for each assay were transferred to a 1.5 ml microcentrifuge tube and centrifuged at 3000 rpm for 5 minutes. The resulting cell pellet was then resuspended in 400 µl of NADH/NAD extracted buffer. The cells were lysed by subjecting them to two rounds of sonication. After sonication, the cells were centrifuged at 13,000 rpm for 10 minutes, and the supernatant was transferred to a different microcentrifuge tube. The samples were then deproteinized using Amicon 10 kDa cut-off spin filters by centrifugation. Subsequently, 50 µl of deproteinized extracted sample from each replicate was transferred into 96-well plates. Into each well, 100 µl of master reaction mixture (comprising 98 µl of NAD cycling buffer and 2 µl of NAD cycling enzyme mix) was added. The plates were then incubated at room temperature for 5 minutes. After the 5-minute incubation with the master reaction mixture, 10 µl of NADH Developer was added to each well. The plates were then incubated at room temperature for 1 to 2 hours. Absorbance at 450 nm was measured to assess the reaction. To measure the NAD level from the bleached cells, cells were directly taken from the bleached plates and mixed into 1 ml of cold Hv-Cas media. The cells were then washed twice. As previously described, the cells for each assay were pelleted in a 1.5 ml microcentrifuge tube at 3000 rpm for 5 minutes. The pellet was then resuspended in 400 µl of NADH/NAD extracted buffer, and the cells were lysed by two rounds of sonication. After sonication, the protein concentration was measured using the Bradford assay. The NAD assay protocol was then followed as described previously. **NADH/NAD assay with HFTV1** NADase activity in *H. gibbonsii* was measured using a commercially available kit (cat. no. [X]). Overnight starter cultures of the empty vector control strain (UG788) and the H-CBASS2-expressing strain (UG784) were grown in Hv-ECas medium at 45°C with shaking. Upon reaching an OD₆₀₀ of 0.8–0.9, cultures were diluted to an OD₆₀₀ of 0.4 in 2 ml of fresh Hv-ECas medium supplemented with 2 mM tryptophan. Cultures were then infected with HFTV1 at an MOI of 0.05. Infected cultures were initially incubated at 45°C for 2 hours, then transferred to 37°C for an additional 5–6 hours. Following incubation, cultures were normalized to equal OD₆₀₀ values, and NADase activity was measured according to the manufacturer’s protocol.

### In vitro evolution experiments

Single isolated colonies of each strain (*H. volcanii* expressing H-CBASS-2 and the virus-infected control strain) were inoculated into Hv-ECas medium supplemented with thymidine and tryptophan and grown to late log or stationary phase. For serial passaging, 100 µL of culture was transferred into 2 mL of fresh medium (1:20 dilution) and allowed to regrow to late log/stationary phase. This process was repeated for approximately 10 passages. Following 3, 6, and 10 passage, 100 µL of each culture was serially diluted in fresh Hv-Cas medium and plated on Hv-Cas agar supplemented with thymidine and tryptophan. Approximately 30–40 individual colonies per sample were screened by PCR to determine the presence of HFPV-1.

### Plaque Assay

For the plaque assay, *H. volcanii* and *H. gibbonsii* strains were grown overnight in Hv-ECas medium and normalized at optical density of 595 nm (OD595). Subsequently, 400 µl of cultures with 0.5M CaCl2 was added to 3-4 ml of 0.2% top-agar (preheated and cooled to 60°C) in Hv-YPC, and the mixture was spread onto rich agar plates with 18% SW. Ten-fold serial dilutions of HFPV-1 and HFTV-1 were prepared in Hv-ECas medium, and 3 µl of each dilution was spotted onto the plates. For HFPV-1, plates were incubated at 30°C for 2 days, after which plaques had formed. Plates were then left at room temperature (∼25°C) for a further 2–3 days to allow plaques to develop and become more distinct, before imaging. For HFTV1, plates were incubated at 37°C for 4–5 days prior to imaging^18^.

### Quantitative real-time PCR (Q-PCR)

To quantify the genome, copy number (gcn) of *H. volcanii* and HFPV-1, we used the CFX Connect Real-Time PCR system (Bio-Rad Laboratories). We collected 1 ml of cultures in biological triplicates at 48 and 72 hours, as well as during the 3rd and 6th passages. These cultures were pelleted at 11,000 × g for 10 minutes at room temperature and supernatant was collected. To assess the amount of secreted viral DNA in the supernatant, DNA was extracted from 200 μl of the supernatant using the Quick-DNA Viral Kit (Zymo Research, D3015). Quantitative PCR (qPCR) was performed using the q-PCRBIO SyGreen Blue mastermix Hi-ROX (Cat. No PB20.16-05) with primers specific to an internal viral protein and a housekeeping gene, the DNA polymerase II small subunit (*polB*). Relative viral abundance was calculated using the ΔΔCt method. Viral Ct values were first normalized to the corresponding *polB* Ct values for each sample (ΔCt), and relative levels were determined by comparison to the indicated control condition. The values presented in the graph therefore represent normalized relative viral abundance rather than raw Ct values.

### RNA extraction and purification

Each strain was grown in Hv-Cas medium at 45°C with shaking to an OD₅₉₅ of ∼0.6, with 2 mM L-tryptophan added where induction was required. Cultures were centrifuged at 6,000 rpm for 5 minutes, and pellets were resuspended in 1 ml Bio-TRI RNA reagent (Bio-Lab). Following addition of 200 µl chloroform, vortexing for 40 seconds, and 3-minute room temperature incubation, samples were centrifuged at 14,000 rpm at 4°C for 15 minutes. RNA was purified using the RNeasy Mini Kit (Qiagen) with on-column DNase I treatment (RNase-Free DNase Set, Qiagen) to remove genomic DNA. RNA was further purified by ethanol/sodium acetate precipitation, dried, quantified by NanoDrop (Thermo Scientific), and stored at −80°C. **RNA preparation for real-time PCR** Following RNA purification, 500 ng of RNA from each sample was reverse-transcribed using random hexamers (Promega) and ImProm-II reverse transcriptase (Promega). cDNA was diluted 1:3 in ultrapure water (Biological Industries) and stored at −20°C. qPCR reactions were performed in a final volume of 20 µl containing 125 ng cDNA, 1× qPCRBIO Fast SyGreen Blue Hi-ROX mix (PCR Biosystems), and 100 nM gene-specific primers. All primers are listed in Table 2. Reactions were run on a CFX Connect Real-Time system (Bio-Rad) under standard cycling conditions, and expression was normalised to the *polB* reference gene.

### Competition assay

To perform head-to-head competition experiments, we used a virus-infected *H. volcanii* strain carrying the H-CBASS-2 system and a virus-infected control strain lacking this system. Each strain was grown overnight to the log phase in Hv-ECas medium supplemented with thymidine and tryptophan. Equal amounts of the log-phase cultures (OD 0.03) were then mixed into fresh Hv-ECas medium and incubated at 45°C for 48 hours with continuous shaking. Samples of 100 µL were taken at 0, 24, and 48 hours, diluted, and plated on Hv-ECas plates containing thymidine and tryptophan. After colonies appeared, the presence of the CBASS gene cluster was verified using internal specific primers for H-CBASS2 by PCR.

### New mutation detection

To check for new mutations and CRISPR spacer acquisition, we selected three individual colonies from each post-infection virus-cured *H-CBASS2*-expressing strain, as well as from the virus-infected wild-type strain after the 10^th^ passage. DNA was extracted using the Blood and Tissue Kit (Qiagen, 69506). Whole genome sequencing was performed by Plasmidsaurus. using Oxford Nanopore Technology with custom analysis and annotation. Mutations in the main chromosome were identified by comparing the main chromsome obtained from each colony to DS2 *H. volcanii* reference genome, using MUMmer 3.0 with default parameters.

### Homology modelling using AlphaFold3

To create a homology model of the H-CBASS2 cyclase, we employed a multi-step process based on AlphaFold3 structure prediction^39^. We first extracted the amino acid sequence of the H-CBASS2 cyclase (accession number: WP_115891644.1) from the NCBI database. The extracted sequence was submitted to the AlphaFold3 server for predicted strcuture generation. After obtaining the model, we conducted a Foldseek^40^ search for structural comparison against the PDB database. The resulting structure was analyzed based on TM-score and RMSD (Root Mean Square Deviation) values. Finally, we used the UCSF Chimera software to visualize the alignments and generate the figures.

### Proteomics

For proteomic analysis, we used a virus-infected *H. volcanii* strain carrying the H-CBASS-2 system and a virus-infected control strain lacking this system. Each strain was grown to the late log phase in Hv-ECas medium. The cultures were centrifuged at 4,500 × g for 45 minutes to collect the cell pellet. The supernatant was collected, and the viruses were precipitated by adding polyethylene glycol (PEG) 6000 to a final concentration of 10% (wt/vol), followed by incubation at 4°C overnight. After overnight incubation, viruses were extracted according to the protocol described by Tomas Alarcon-Schumacher et al. ^16^. The cell pellet and extracted viral particles were sent directly to the Smoler Proteomics Center at the Lorry I. Lokey Interdisciplinary Center for Life Sciences and Engineering, Technion, Israel. The samples were digested with trypsin and analyzed by LC-MS/MS using the Q Exactive HFX mass spectrometer. Data analysis was performed using Proteome Discoverer 2.4 software and the Sequest search engine against both a specific database and a decoy database to determine the false discovery rate (FDR). All identified peptides were filtered with a high-confidence 1% FDR threshold. Quantification was carried out by calculating the peak area of each peptide, with protein abundance represented by the sum of all associated peptide group abundances. **Host protein analysis** The mean raw peptide count was calculated per experimental condition. The difference in mean protein abundance between control cells and H-CBASS2-expressing cells (Δ control − H-CBASS2), as well as between virus-infected control cells and virus-infected H-CBASS2-expressing cells (Δ control+ virus− H-CBASS2+virus), was calculated for each protein feature. While Δ control − H-CBASS2 was normally distributed around zero, Δ control+virus−H-CBASS2+virus was heavily skewed to the right, indicating differential protein abundance upon viral infection in H-CBASS2-expressing cells. Protein features with Δ control+virus−H-CBASS2+virus values of less than −4 or greater than 22 were selected for display in the heatmap.

## Author contributions

U.G. and D.K.C. conceived and designed the study. D.K.C., H.S, and D. V, performed the experiments. D.K.C., H. S, N.G., and L.R. analyzed the data. A.B.I, and M.P generated transposon insertion mutant. D.K.C. prepared the figures and analyzed the data with input from U.G. The manuscript was written by D.K.C., with U.G. and L.R. contributing to its editing. All authors read and approved the final draft.

## Acknowledgements

The authors thank Prof. Susanne Erdmann for providing HFPV-1 and Dr. Tessa E F Quax for providing HFTV1, Sharon Navok for assistance with the development and execution of plaque assays, and Alex Barbul for help with confocal microscopy. The authors also thank Dr. Tomas Alarcón-Schumacher, Dr. Israela Turgeman-Grott and Neta Altman-Price for helpful discussions. The work is dedicated to the memory of Rachel Schreiber.

