## Extended figures for "An archaeal CBASS shows protective activity against both chronic and lytic viruses"

### An archaeal CBASS system eliminates viruses without killing the host cells

```
Query  3    ELPSKFEEFLSNIRPTDKQISDYVDGHEQLRERLSNDGDLSEIHVSDFLQGSYARRTAVK  62
            EL  +F EFL+NIRPTD Q   D+   G   LRERL N   L EI VS FLQGS  R TA++
Sbjct  3    ELQPQFNEFLANIRPTDTQKEDWKSGARTLRERLKNFEPLKEIVVSTFLQGSIRRSTAIR  62

Query  63   PIGDEKSDVDIVFVTNLPKSEYTASDAMELCEPFLNRYYPGQWEPNQRSYKIELNKVEMD  122
            P+GD++ DVDIV VTNL   +   + +DAM+L  PFL +YYPG+WE   RS+ I L+ VE+D
Sbjct  63   PLGDKRPDVDIVVVTNLDHTRMSPTDAMDLFIPFLEKYYPGKWETQGRSFGITLSYVELD  122

Query  123  LVLTAAPSEAV----INELSAPGSIGKANIANIADQQDMGIIAKSIDNTFEGAEKN----  174
            LV+TA P         + +L    S+   N  ++ +Q D  +       NT   +E N
Sbjct  123  LVITAIPESGAEKSHLEQLYKSESVLTVN--SLEEQTDWRLNKSWTPNTGWLSESNSAQV  180

Query  175  -------WQDEPLDIPDRELDSWDQTHPLATLDWTQNKNDRTNGHYINVVKALKWWRRTQ  227
                   W+   PL +PDRE + W +THPLA + WT  KN   NGHYIN+V+A+KWWR+
Sbjct  181  EDAPASEWKAHPLVLPDREKNEWGRTHPLAQIRWTAEKNRLCNGHYINLVRAVKWWRQQN  240

Query  228  V-DFPERPKSYPLERLIGECCPDYISSVAEGVVRAFDVFIEKYESNAEHEDTPVLGQHGI  286
              D P+ PK YPLE LIG     +  +S+A+G+V+  D F+ ++ +     +  P L  HG+
Sbjct  241  SEDLPKYPKGYPLEHLIGNALDNGTTSMAQGLVQLMDTFLSRWAAIYNQKSKPWLSDHGV  300

Query  287  PENNVLARLEGRDFAAFYGRVTDASEVAQRAYEEEDKEQSALYWREMFGEEFPLIGDSDD  346
             E++V+ARL    DF  +FY   +  A+E+A+  A    E+  ++SA  WR++FG +FPL G
Sbjct  301  AEHDVMARLTAEDFCSFYEGIASAAEIARNALASEEPQESAQLWRQLFGSKFPLPGPQGG  360

Query  347  DDTDDESTATFTPPNNTA  364
            D          FT P+  A
Sbjct  361  D-----RNGGFTTPSKPA  373
```

**Extended Fig.1 A pairwise BLASTP alignment of the H-CBASS2 cyclase with the *E. cloacae* cyclase (CdnD).** The catalytically important residue, highlighted in yellow, has been mutated to render the cyclase inactive. The query represents the H-CBASS2 cyclase and the subject represents the *E. cloacae* cyclase (CdnD).

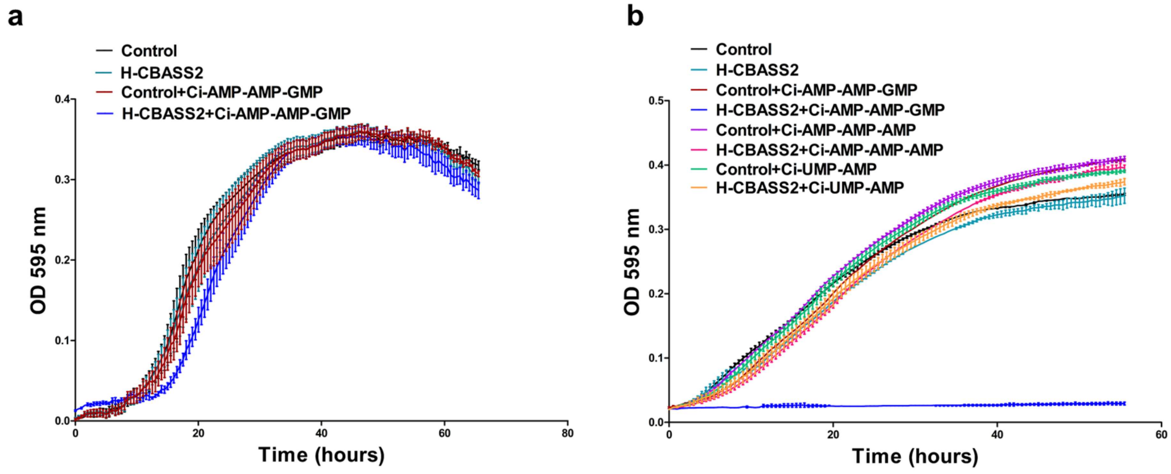


**Extended Fig. 2. Exogenous cyclic-AMP-AMP-GMP activates H-CBASS2 in the absence of virus. (a)** Growth analysis of *H. volcanii* strains expressing H-CBASS2 (blue) and the empty vector control (red) in the presence of exogenous cyclic-AMP-AMP-GMP at a lower concentration (6 µM). Untreated empty vector control and untreated H-CBASS2 strains are shown in black and cyan, respectively. Lines represent the mean of at least 3 biological replicates, each with two technical replicates. **(b)** Growth analysis of *H. volcanii* strains expressing H-CBASS2 in the presence of different cyclic signaling molecules at a higher concentration (12 µM). In the presence of exogenous cyclic-AMP-AMP-GMP, H-CBASS2-expressing and empty vector control strains are shown in blue and red, respectively. Untreated H-CBASS2-expressing and untreated empty vector control strains are shown in cyan and black, respectively. In the presence of cyclic-AMP-AMP-AMP, H-CBASS2-expressing and empty vector control strains are shown in pink and purple, respectively. In the presence of cyclic-UMP-AMP, H-CBASS2-expressing and empty vector control strains are shown in brown and green, respectively. Lines represent the mean of at least 2 biological replicates, each with two technical replicates.

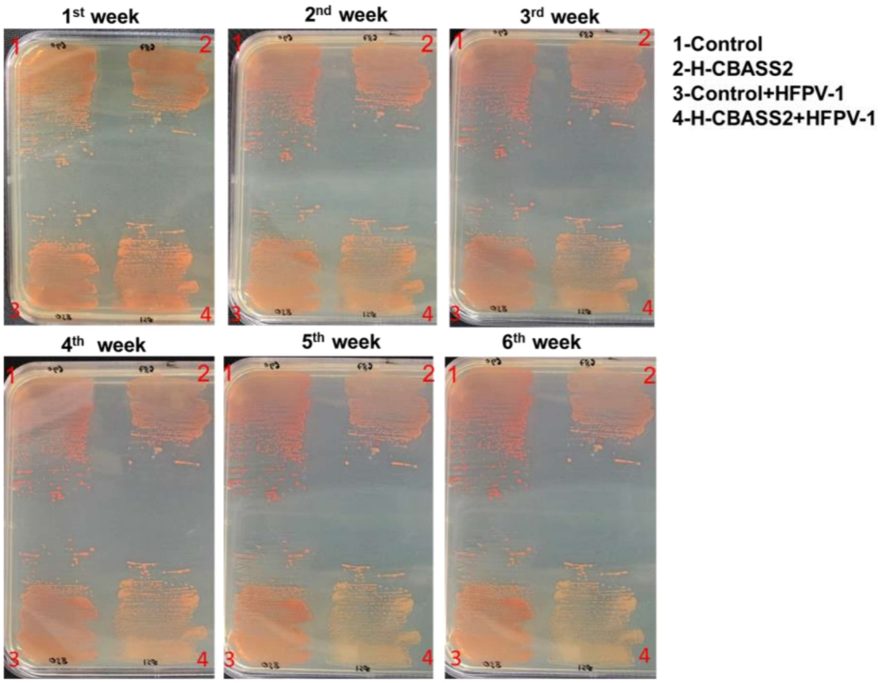


**Extended Fig.3 Representative images showing a gradual bleaching of the virus-infected H-CBASS2-expressing colonies.** A time series analysis was conducted on colony color with or without viral infection in the H-CBASS2-expressing strain, as well as in the infected control containing an empty vector.

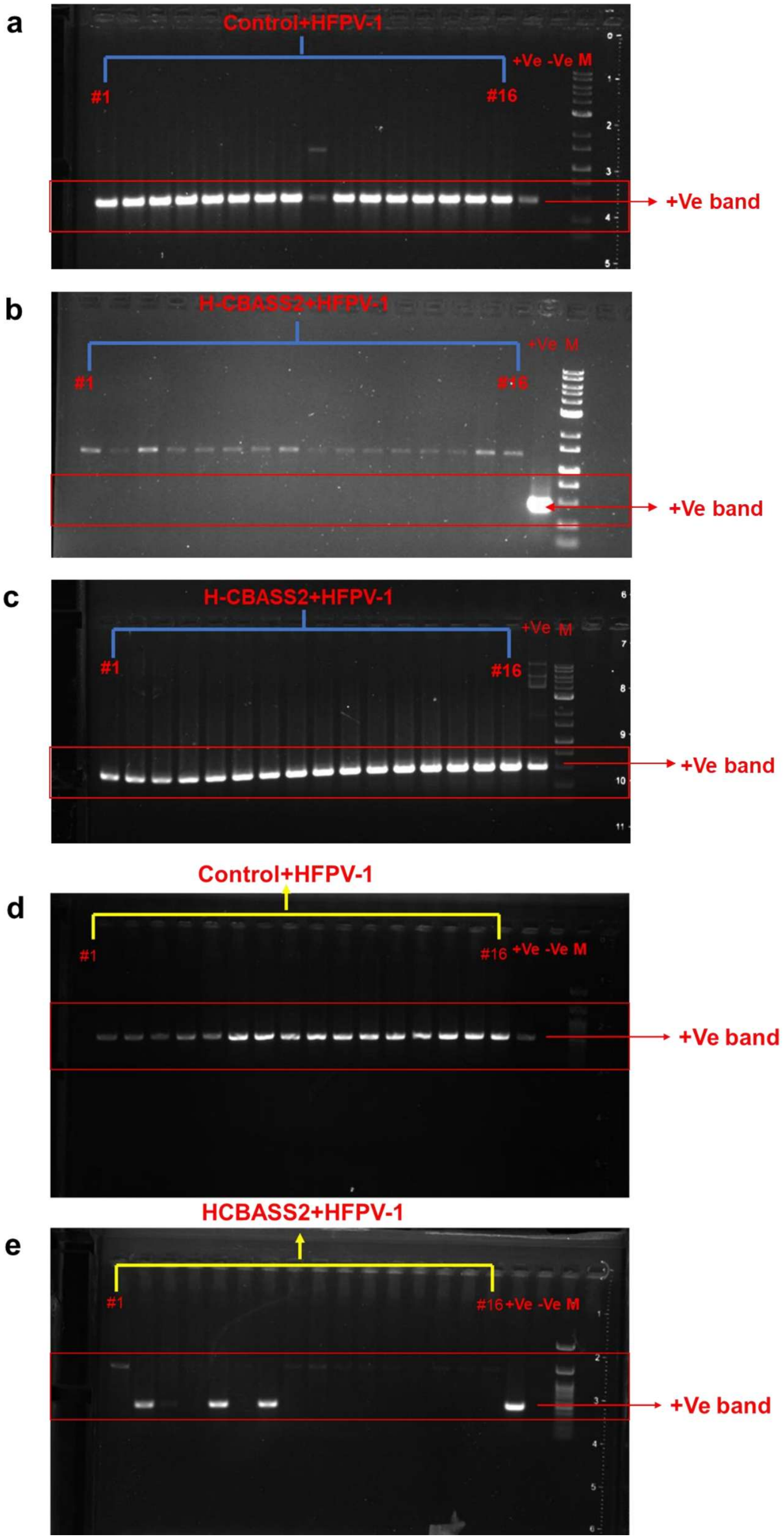


**Extended Fig.4 Most H-CBASS2-expressing colonies are virus-negative after passage 10.** The results of ethidium bromide-stained agarose gel electrophoresis of products from colony PCR. **a**) Virus detection via PCR after the 10$^{th}$ passage in virus-infected *H. volcanii* strains containing an empty vector (control). The gel displays colonies #1 to 16 from left to right after the 10th passage, along with positive control, negative control, and a 1Kb DNA ladder (NEB). The expected band size is ~500 bp. **b**) Virus detection via PCR after the 10$^{th}$ passage in virus-infected *H. volcanii* strains expressing H-CBASS2. The gel displays colonies #1 to 16 from left to right, with a positive control and a 1Kb DNA ladder (NEB). The expected band size is ~500 bp. Band display ~1500 bp is nonspecific. **c**) H-CBASS2 detection via PCR by amplifying an internal region after the 10$^{th}$ passage in virus-infected *H. volcanii* strains expressing H-CBASS2. The gel shows colonies #1 to 16 from left to right, with a positive control and a 1Kb DNA ladder (NEB). The expected band size is ~550 bp. **d**) Virus detection via PCR after the 3$^{rd}$ passage in virus-infected *H. volcanii* strains containing an empty vector (control). The gel displays colonies #1 to 16 from left to right after the 10th passage, along with positive control, negative control, and a 1Kb DNA ladder (NEB). The expected band size is ~500 bp. **e**) Virus detection via PCR after the 3$^{rd}$ passage in virus-infected *H. volcanii* strains expressing H-CBASS2. The gel displays colonies #1 to 16 from left to right, with a positive control and a 1Kb DNA ladder (NEB). The expected band size is ~500 bp. Band display ~1500 bp is nonspecific.

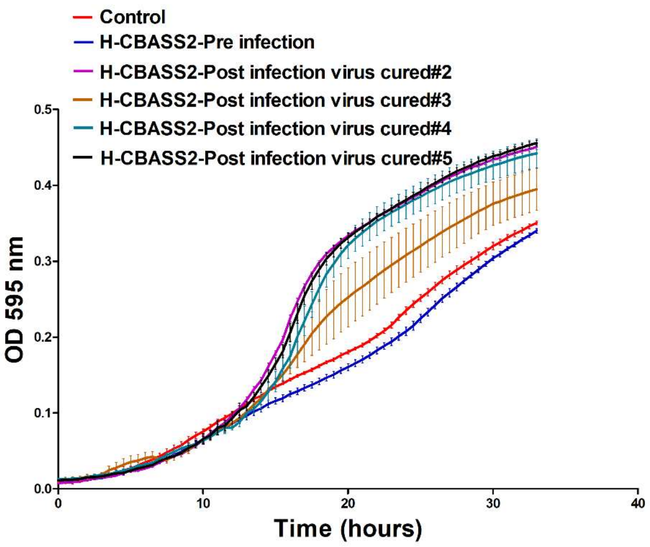


**Extended Fig.5. Virus-cured H-CBASS2–expressing colonies exhibit faster growth.** Growth curves of individual virus-cured (post-infection) H-CBASS2–expressing strains are shown. Post-infection strains derived from virus-cured colonies #2, 3, 4, and 5 are shown (purple, brown, cyan, and black), along with the pre-infection H-CBASS2–expressing strain (blue) and a non-infected control strain carrying an empty vector (red). Lines represent the mean of at least two biological replicates, each with three technical replicates.

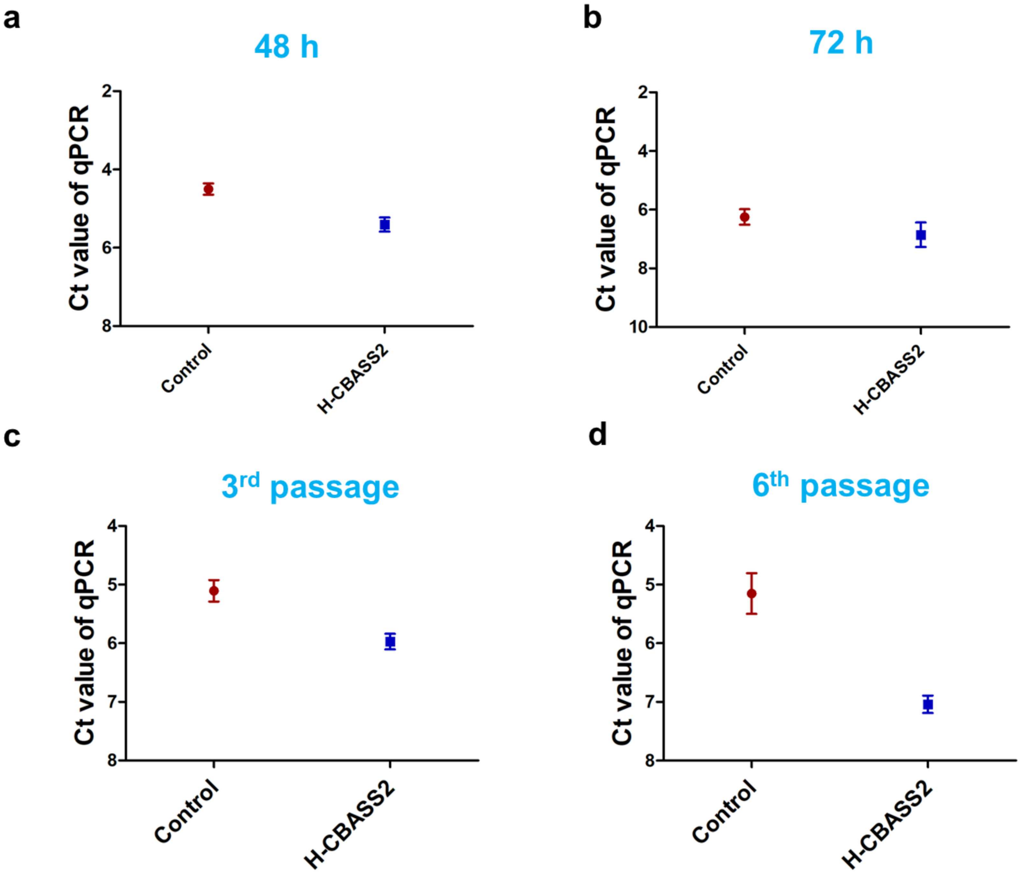


**Extended Fig.6- CBASS-expressing cells exhibit a reduction in viral DNA levels.** qPCR was performed to quantify the amount of viral DNA in the supernatant of HFPV-1 infected *H. volcanii* cells after 48 and 72 hours of growth, as well as after the 3rd and 6th passages, using specific primers targeting the HFPV-1 genome. **a**) Ct values for free viral DNA in the supernatant after 48 hours. **b**) Ct values for free viral DNA in the supernatant after 72 hours. **c**) Ct values for free viral DNA in the supernatant after the 3rd passage. **d**) Ct values for free viral DNA in the supernatant after the 6th passage.

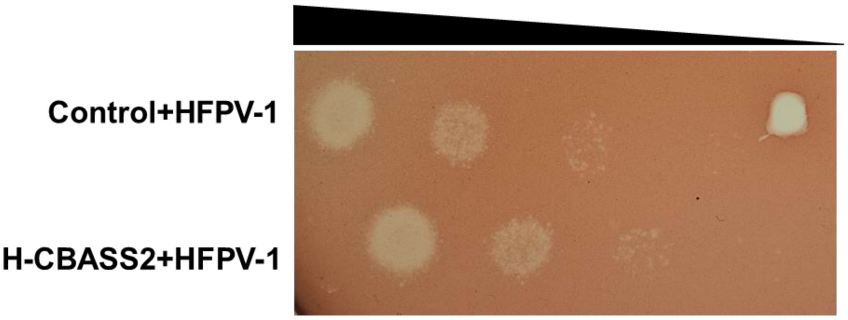

**Extended Fig. 7. Infection assay using HFPV-1 extracted from HFPV-1-infected strains expressing H-CBASS2 and control cells.** To determine whether H-CBASS2 affects viral egress, we purified HFPV-1 from the supernatant of both H-CBASS2-expressing and empty vector control cells after 7 days of continuous growth. Plaque assays were subsequently performed using the purified virus on the same *H. volcanii* UG690 reporter strain. Images are representative of at least three independent biological replicates.

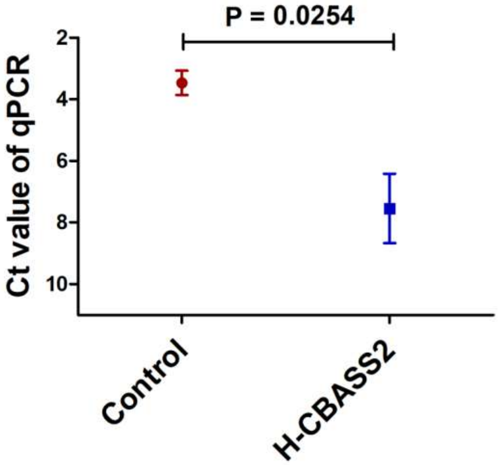


**Extended Fig. 8. H-CBASS2 expression decreases the levels of a putative transcriptional regulator.** H-CBASS2-expressing cells exhibit reduced viral copy numbers. qRT-PCR was performed on exponentially growing *H. volcanii* cells infected with HFPV-1, comparing H-CBASS2-expressing cells with control cells carrying an empty vector. The resulting Ct values are shown in the graph. Data represent at least three independent experiments. Statistical significance was assessed using a paired-sample t-test.

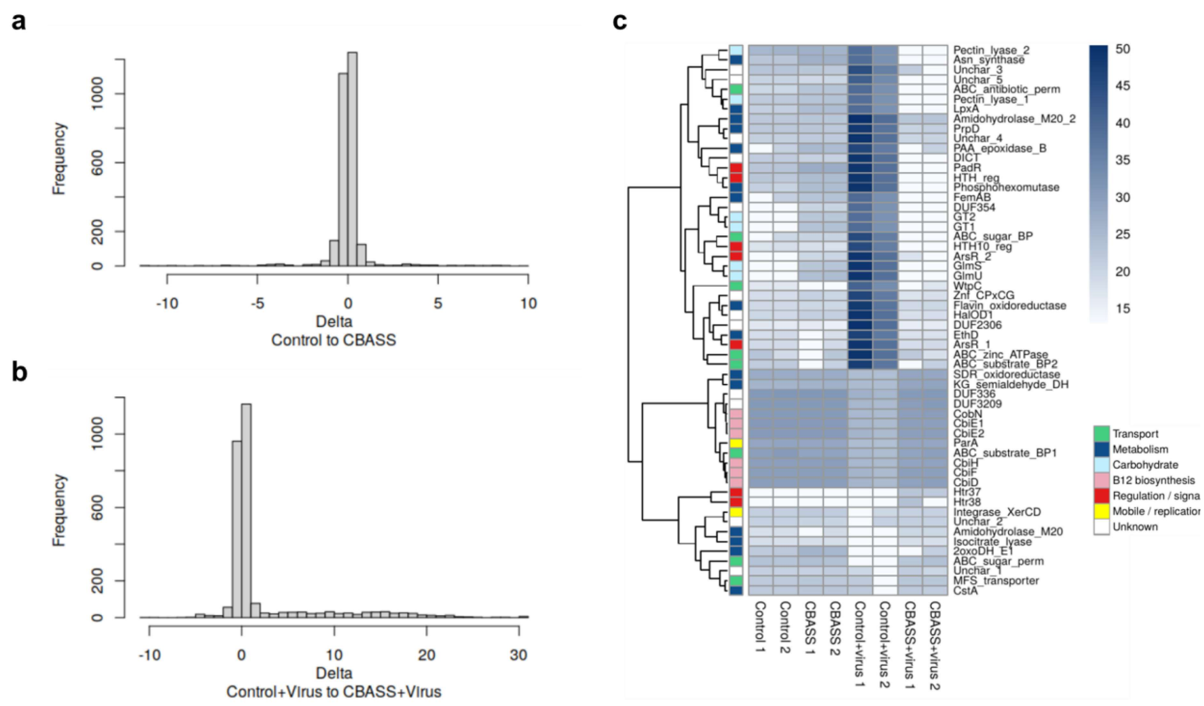


**Figure 9. H-CBASS2 expression affects host proteome during HFPV-1 infection.** a & b) The mean raw peptide count was calculated per experimental condition. The difference in mean protein abundance between control cells and H-CBASS2-expressing cells (Δ control − H-CBASS2), as well as between virus-infected control cells and virus-infected H-CBASS2-expressing cells (Δ control+ virus− H-CBASS2+virus), was calculated for each protein feature. While Δ control − H-CBASS2 was normally distributed around zero, Δ control+virus−H-CBASS2+virus was heavily skewed to the right, indicating differential protein abundance upon viral infection in H-CBASS2-expressing cells. **c**) Heatmap of differential host protein abundance in H-CBASS2-expressing *H. volcanii* cells in the presence or absence of HFPV-1 infection. Each row represents an individual protein, and colour intensity reflects relative protein abundance levels. Protein features with Δ control+virus−H-CBASS2+virus values of less than −4 or greater than 22 were selected for display in the heatmap.

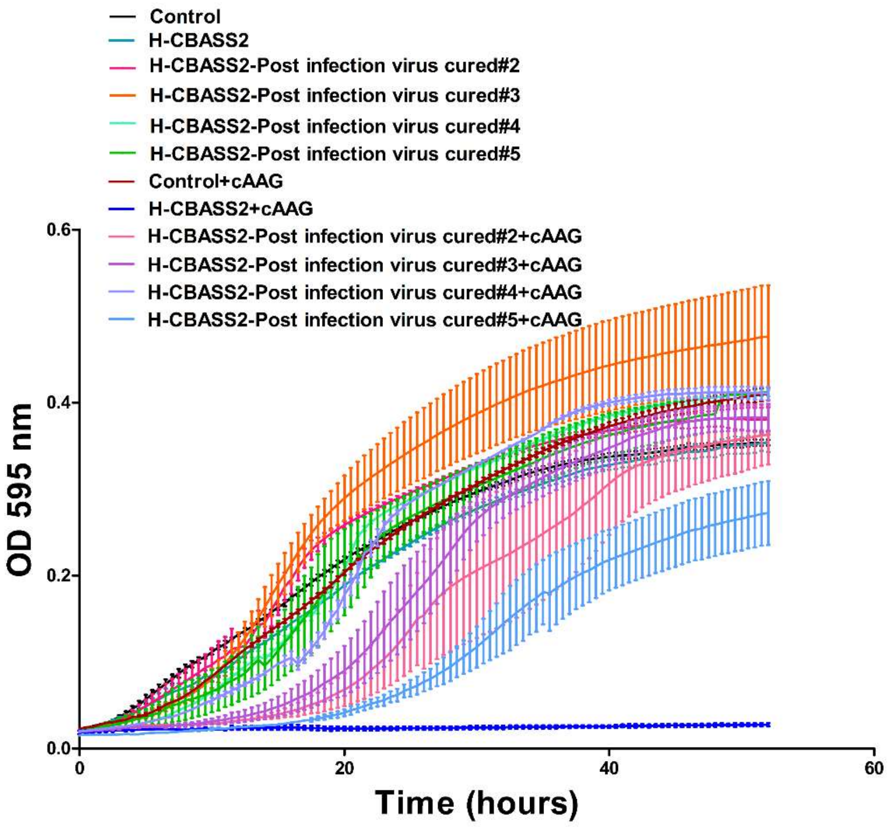


**Extended Fig. 10. Growth analysis of virus-cured *H. volcanii* strains expressing H-CBASS2 in the presence of exogenous cyclic-AMP-AMP-GMP (cAAG).** Growth curves of *H. volcanii* strains expressing H-CBASS2 and empty vector control strains, along with four independently derived virus-cured isolates (clones #2–#5). Upon exposure to 12 µM exogenous cAAG, virus-cured strains displayed a variable resistance phenotype compared to the parental H-CBASS2-expressing strain, which exhibited near-complete growth inhibition in the presence of cAAG. The untreated H-CBASS2-expressing strain (cyan) and empty vector control (black) are shown for reference. Lines represent the mean ± standard deviation of at least three biological replicates, each with two technical replicates.
